# Scc2 counteracts a Wapl-independent mechanism that releases cohesin from chromosomes during G1 but is unnecessary during S phase for establishing cohesion

**DOI:** 10.1101/513960

**Authors:** Madhusudhan Srinivasan, Naomi J. Petela, Johanna C. Scheinost, James Collier, Menelaos Voulgaris, Maurici Brunet-Roig, Frederic Beckouët, Bin Hu, Kim A. Nasmyth

## Abstract

Cohesin’s association with chromosomes is determined by loading dependent on the Scc2/4 complex and release due to Wapl. We show here that Scc2/4 is not merely a loading complex and that it actively maintains cohesin on chromosomes during G1. It does so by blocking a Wapl-independent release reaction that requires opening the cohesin ring at its Smc3/Scc1 interface as well as the D loop of Smc1’s ATPase. The Wapl-independent release mechanism is switched off as cells activate Cdk1 and enter G2/M and cannot be turned back on without cohesin’s dissociation from chromosomes. The latter phenomenon enabled us to show that cohesin that has already captured DNA in a Scc2-dependent manner before replication no longer requires Scc2 to capture sister DNAs during S phase.

## Introduction

Accurate chromosome segregation is an essential aspect of cell proliferation. This remarkable feat is only possible because monumental topological problems posed by the sheer size and physical properties of DNA are overcome by highly conserved DNA motors, namely condensin and cohesin. Replicated DNA is weaved into discrete chromatids during mitosis by condensin (Hirano et al., 1997) while sister chromatids are held together by cohesin, which is essential for their bi-orientation on mitotic spindles.

Both complexes contain a pair of rod-shaped Smc proteins (Smc1/3 in cohesin) whose association via their hinge domains creates V-shaped heterodimers with ATPase domains at their vertices. These are interconnected by kleisin subunits to form trimeric rings, whose activity is regulated by a set of hook-shaped proteins composed of HEAT repeats known as HAWKs (HEAT repeat proteins Associated With Kleisins) (Wells et al., 2017). Regulation by HAWKs distinguishes cohesin and condensin from bacterial Smc/kleisin complexes and the eukaryotic Smc5/6 complex, whose kleisin subunits associate instead with tandem winged helical domain proteins called KITEs (Palecek and Gruber, 2015). Cohesin has three HAWKs: Scc3 is permanently bound while Scc2/Nipbl and Pds5 appear interchangeable (Petela et al., 2018).

Condensin has the remarkable ability to form and expand in a processive manner DNA loops in vitro (Ganji et al., 2018), an activity known as loop extrusion (LE) previously postulated to explain how condensin transforms interphase chromosomes into thread-like chromatids while at the same time accumulating along their longitudinal axis (Goloborodko et al., 2016; Nasmyth, 2001; Naumova et al., 2013). Cohesin has more diverse activities. In addition to its canonical role of holding together sister chromatids, cohesin also organizes interphase chromatin into defined territories called TADs (Topologically Associated Domains), a process also thought to be driven by loop extrusion (Fudenberg et al., 2016; Rao et al., 2017). It has been proposed that regulation of cohesin’s loop extrusion by the site-specific DNA binding protein CTCF underlies how the latter insulates certain enhancers from non-cognate promoters at defined stages of development. Cohesin is thought to mediate cohesion by entrapping sister DNAs inside its tripartite ring (Srinivasan et al., 2018). It can also entrap individual DNAs prior to DNA replication and this may be a feature of its chromosomal association throughout the cell cycle. Whether DNAs are entrapped within cohesin rings during LE is not known.

Loading of cohesin onto chromosomes as well as entrapment of mini-chromosome DNAs by cohesin rings depends on both Scc3 and Scc2 but not on Pds5. Loading also requires Scc4, which binds to an unstructured N-terminal domain within Scc2. Because neither Scc2 nor Scc4 are required to maintain cohesion following S phase, Scc2/4 has long been thought to function merely as a “loading complex”(Ciosk et al., 2000). However, the finding that Scc2 associates with chromosomal cohesin long after loading (Rhodes et al., 2017a) suggests that this may not be the whole story. Because it is essential for activating cohesin’s ATPase (Petela et al., 2018), Scc2 may have a key role in activating the DNA translocase activity that powers LE.

DNAs are released from cohesin rings by two mechanisms, either through kleisin cleavage by separase, which occurs at the onset of anaphase (Uhlmann et al., 1999), or at other stages of the cell cycle via a separase-independent mechanism that involves the binding to Pds5 and Scc3 of a fourth regulatory subunit Wapl. This induces in an ATP-dependent manner disengagement of the ring’s Smc3/kleisin interface, thereby creating a gate through which DNAs can escape (Beckouet et al., 2016; Murayama and Uhlmann, 2015). The steady state level of chromatin associated cohesin during G1 is therefore determined by the rates of Scc2 catalyzed loading and Wapl-dependent releasing activity (RA). Because it would destroy sister chromatid cohesion, Wapl-dependent cohesin release is neutralized during DNA replication through acetylation of Smc3 K112 and K113 by Eco1 (Ben-Shahar et al., 2008; Unal et al., 2008). Though normally essential for cell viability, Eco1 is dispensable in mutants defective in release (Chan et al., 2012; Rowland et al., 2009; Srinivasan et al., 2018).

Both cohesion establishment and the Smc3 acetylation needed to maintain it are tightly coupled to DNA replication. For example, cohesin that loads onto chromosomes during G2 cannot connect sister DNAs (Haering et al., 2004) and mutation of the non-essential fork-associated proteins (Ctf4, Ctf18, Tof1, Csm3, Mrc1and Chl1) causes cohesion defects without adversely affecting replication (Borges et al., 2013; Zheng et al., 2018). Nevertheless, the mechanism by which cohesion is established remains poorly understood. Photo-bleaching experiments showing that replication fork passage does not per se induce cohesin’s dissociation (Rhodes et al., 2017b) suggest either that replication forks actually pass through cohesin rings that have entrapped DNA ahead of the fork or that these cohesin rings are opened transiently as they are passed on to, and subsequently entrap sisters.

One way of providing insight into cohesion establishment would be to determine whether establishment depends on Scc2. Replication through rings should not require a second Scc2-dependent loading reaction while ring opening and reloading on lagging or leading (or both) strands would be expected to do so. The recent observation that cohesin loaded onto a double stranded DNA in vitro is capable of capturing a second single stranded DNA molecule in a manner dependent on Scc2 (Murayama et al., 2018) raises the possibility that cohesion is established by rings associated with a leading strand capturing the lagging strand through Scc2-catalysed ring opening. If so, Scc2 must be required during replication itself as well for loading cohesin onto un-replicated chromatin.

We therefore set out to answer the following simple question: Is Scc2 required to build cohesion during S phase in cells that have already loaded cohesin onto chromosomes during G1? Because Scc2 is required during G1 to reload cohesin that has been removed from chromosomes, it was necessary to perform our experiment in cells lacking RA. Unexpectedly, inactivation of Scc2 in pre-replicative cells causes cohesin unloading throughout the genome, even in cells where turnover has been abrogated by mutations (e.g *wplΔ*) that normally abolish Wapl-dependent RA. We discovered that release in G1 cells involving disengagement of the Smc3/Scc1 interface is in fact a Wapl-independent process that is actively blocked by Scc2. Scc2 is therefore not merely a loader and Wapl is not an intrinsic aspect of RA. Wapl-independent RA is switched off as cells undergo S phase in a manner that does not require either replication or acetylation of Smc3 by Eco1 but involves Cdk1. Because it cannot be switched back on (upon Cdk1’s subsequent inactivation) without cohesin’s removal from chromosomes, we were able to show that cohesin loaded prior to replication can create cohesion without Scc2, a finding that has profound implications regarding the mechanism of cohesion establishment.

## Results

### A Wapl-independent activity releases cohesin from chromosomes in G1 cells

To measure DNA entrapment within cohesin rings, we used cells containing cysteine pairs at all of cohesin ring’s three interfaces (6C) that can be cross-linked by the homobifunctional crosslinker BMOE (Gligoris et al., 2014). Following SDS treatment, gel electrophoresis and Southern blotting reveals two types of circular mini-chromosomes associated with 6C cohesin only when cells are treated with BMOE: catanated monomers (CMs) that correspond to monomeric supercoiled DNAs catenated by a single cohesin ring and catanated dimers (CDs) that correspond to sister DNAs catenated by a single ring (Fig. 1A)(Srinivasan et al., 2018). CMs are formed when cohesin loads onto chromosomes either before or after (see below) DNA replication while CDs only form when cohesin builds cohesion during S phase.

**Figure 1.**
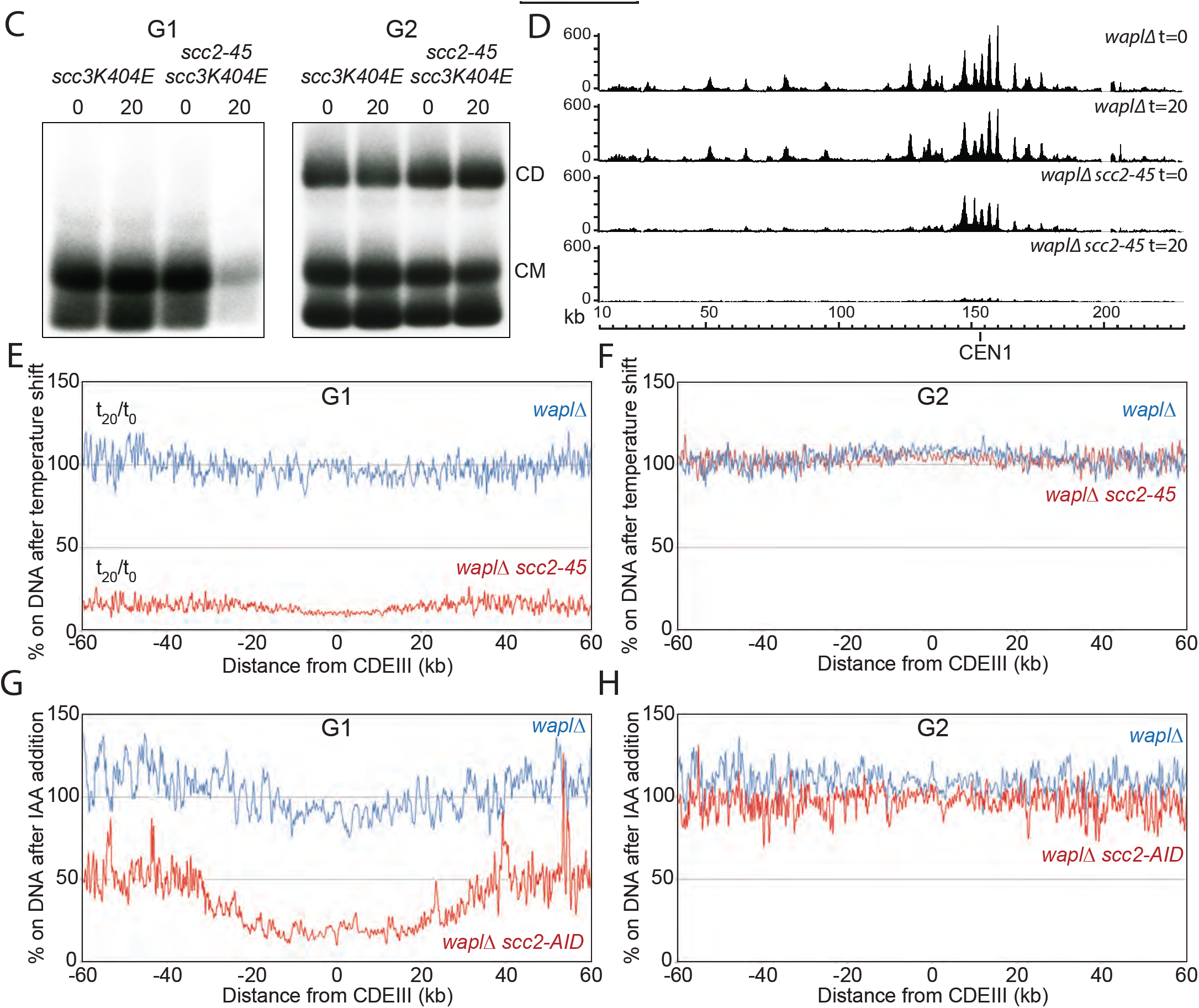
A Wapl-independent activity releases cohesin from chromosomes in G1 cells. **(A)** Schematic of the mini-chromosome IP assay. 6C strain (K23889) with cysteine pairs at all three ring subunit interfaces (2C Smc3: E570C S1043C, 2C Smc1: G22C K639C and 2C Scc1 C56 A547C) and carrying a 2.3 kb circular mini-chromosome were subjected to in vivo crosslinking with BMOE. DNAs associated with cohesin immune-precipitates (Scc1-PK6) were denatured with SDS and separated by agarose gel electrophoresis. Southern blotting reveals two forms of DNA unique to cells treated with BMOE: CMs and CDs. **(B)** WT (K3972) and *scc2-45* (K25238) 6C strains expressing galactose-inducible nondegradable sic1 were arrested in late G1 at 25°C as described in STAR methods. The cultures were shifted to 37°C for 20 minutes, aliquots drawn before (0) and after (20) temperature shift were subjected to mini-chromosome IP. **(C)** *scc3K404E* (K25313) and *scc3K404E scc2-45* (K25316) 6C strains were either arrested in late G1 like described in (B) or in G2 with nocodazole at 25°C and subjected to mini-chromosome IP like described in (B) **(D)** *waplΔ* (K22296) and *waplΔ scc2-45* (K22294) strains were arrested in late G1 at 25°C and subjected to temperature shift to 37°C for 20 minutes. 0 and 20 minute samples were analyzed by calibrated ChIP-sequencing (Scc1-PK6). ChIP profiles along chromosomes I is shown. **(E)** Data form (D) is plotted to show the ratio of average cohesin levels 60 kb on either side of all 16 centromeres before and 20 minutes after the temperature shift. **(F)** the ratio of average cohesin levels 60 kb on either side of all 16 centromeres before and 20 minutes after the temperature shift of *waplΔ* (K22296) and *waplΔ scc2-45* (K22294) strains were arrested in G2 at 25°C. **(G and H)** *waplΔ* (K20891) and *scc2-3XmAID waplΔ* (K26831) were arrested in either late G1 or G2 and treated with auxin (IAA) for 60 minutes and subjected to Cal-ChIP-Seq. ratio of average cohesin levels before and after IAA addition is plotted. See Fig S1 for supporting data.

To ascertain whether cohesin’s “loading complex” is required to establish cohesion during S phase as well as to load cohesin onto chromosomes in the first place, we set out to determine whether its Scc2 subunit is required to convert CMs formed during G1 into CDs during replication. To do this, we first established a protocol for inactivating Scc2 in G1 cells that accumulate Scc1 to high levels and load cohesin onto chromosomes. The simplest way of arresting yeast cells in G1, namely incubating *MATa* cells with *α* factor pheromone, is not appropriate because Scc1 cleavage that commences during anaphase persists during pheromone induced G1 arrest. Moreover, there is very little de novo Scc1 synthesis at this stage of the cell cycle. We therefore arrested wild type (WT) and *scc2-45* (a temperature sensitive *SCC2* allele) 6C cells in late G1 by expression of a non-degradable form of the CDK1 inhibitor Sic1 at the permissive temperature (25°C). Under these conditions, activation of transcription by SBF and MBF turns on not only Scc1 synthesis but also that of securin, which inactivates separase. Crucially, the cohesin that accumulates forms CMs (Srinivasan et al., 2018), which disappear in *scc2-45* but not WT when cells are shifted to 37°C for 20 min (Fig. 1B).

The simplest explanation for cohesin’s dissociation from mini-chromosomes is that Scc2 is required to re-load chromosomal cohesin that is continually turning over due to Wapl-mediated RA, which is active in these cells (Chan et al., 2012; Lopez-Serra et al., 2013). To test this, we repeated the experiment using cells carrying *scc3K404E* (Fig. 1C & S1A) or *pds5S81R* (Fig. S1B) mutations that abolish RA in otherwise wild type cells (Beckouet et al., 2016). Surprisingly, neither mutation abrogated loss of CMs in *scc2-45* cells (Fig. 1C and S1B). Calibrated ChIP-seq confirmed that cohesin dissociates from the entire genome, not merely from *CEN* proximal sequences present in mini-chromosomes (Fig. S1C).

It is possible that neither *pds5S81R* nor *scc3K404E* completely abolish Wapl-dependent RA. We therefore tried to delete *WPL1*, but discovered that this adversely affects proliferation of *scc2-45* cells at 25°C. Fortunately, this synthetic lethality is relieved by a point mutation in *SMC3* (*smc3R1008I*) that has little or no phenotype on its own and importantly does not restore RA as it enhances not reduces proliferation of *wpl1Δ eco1Δ* (Fig. S1D). The mechanism by which *smc3R1008I* works will be presented elsewhere. The mutation was used in all further experiments involving *wpl1Δ scc2-45* cells.

Calibrated ChIP-seq showed that shifting Sic1-arrested cells to 37°C for 20 min had no effect on chromosomal cohesin in *smc3R1008I wpl1Δ SCC2* cells but caused a major reduction throughout the genome in *smc3R1008I wpl1Δ scc2-45* cells (Fig. 1D & S1E). Fig. 1E documents this effect by plotting the ratio of average cohesin levels 60 kb either side of all 16 centromeres before and 20 minutes after the temperature shift (Fig. 1E). The key point is that the −60 kb to +60 kb ratio profile from *SCC2* (blue) is clearly separated from that from *scc2-45* (red) cells. Overall, inactivation of *scc-45* causes about a sevenfold reduction in chromosomal cohesin.

Remarkably, inactivation of *scc2-45* had no effect on chromosomal cohesin in cells arrested in G2/M phase by nocodazole. Thus, neither CMs nor CDs (Fig. 1C) were altered by shifting *scc3K404E scc2-45* cells to 37°C. Likewise, the calibrated ChIP-seq −60 kb to +60 kb temperature shift ratio profiles of *smc3R1008I wpl1Δ SCC2* (blue) and *smc3R1008I wpl1Δ scc2-45* (red) were indistinguishable (Fig. 1F & S1F).

To ensure that this effect is not specific to *scc2-45*, we repeated the experiment with an Auxin degron allele (*SCC2-3XmAID*). Plotting the −60 kb to +60 kb ratio profiles before and after addition of synthetic auxin (Indole-3-acetic acid) for 60 min again revealed a major discrepancy between *SCC2* (blue) and *SCC2-3XmAID* (red) in G1 (Fig. 1G) but not in G2 (Fig. 1H) cells. Thus, Scc2 depletion also causes cohesin’s dissociation from G1 but not G2 chromatin genome wide.

Because Scc2’s inactivation is not accompanied by any change in the overall levels of Scc1 (Fig. S1G), we conclude that Scc2 is essential to maintain cohesin’s association with chromosomes during late G1 even when there is no turnover. This implies that G1 but not G2 cohesin has the ability to dissociate from chromatin through a Wapl-independent mechanism and that Scc2 is required to counteract this activity. Three important corollaries follow. First, Scc2 is not merely involved in loading cohesin onto chromosomes but acts long afterwards to prevent release. Second, contrary to prevailing wisdom, Wapl is in fact not essential for cohesin to dissociate from chromosomes. Third, this Wapl-independent release mechanism is cell cycle regulated and turned off in G2 cells, with the result that Scc2 is no longer required to maintain cohesin on chromosomes after replication. The notion that Scc2 is active on chromosomal cohesin is consistent with the finding that Scc2/Nipbl associates transiently with chromosomal cohesin in mammalian cells even when that latter does not turnover (Rhodes et al., 2017a).

### Scc2 does not require Scc4 to block Wapl-independent release

How does Scc2 block release? Our first step was to investigate Scc4’s role. Calibrated ChIP-seq showed that the temperature shift profiles of *wpl1Δ SCC4* cells and *wpl1Δ scc4-4* cells arrested in late G1 by Sic1 are very similar if not identical (Fig. 2A). Thus, in contrast to Scc2, inactivation of Scc4 in late G1 has little or no effect on chromosomal cohesin. Because Scc4 is required for de novo loading (Fig S2A) (Petela et al., 2018), this observation confirms that there is indeed little or no turnover of chromosomal cohesin in late G1 cells lacking Wapl, which is consistent with imaging studies in both in yeast (Chan et al., 2012) and mammalian cells (Rhodes et al., 2017a). Scc2’s association with Scc4 does not appear necessary to block release.

**Figure 2.**
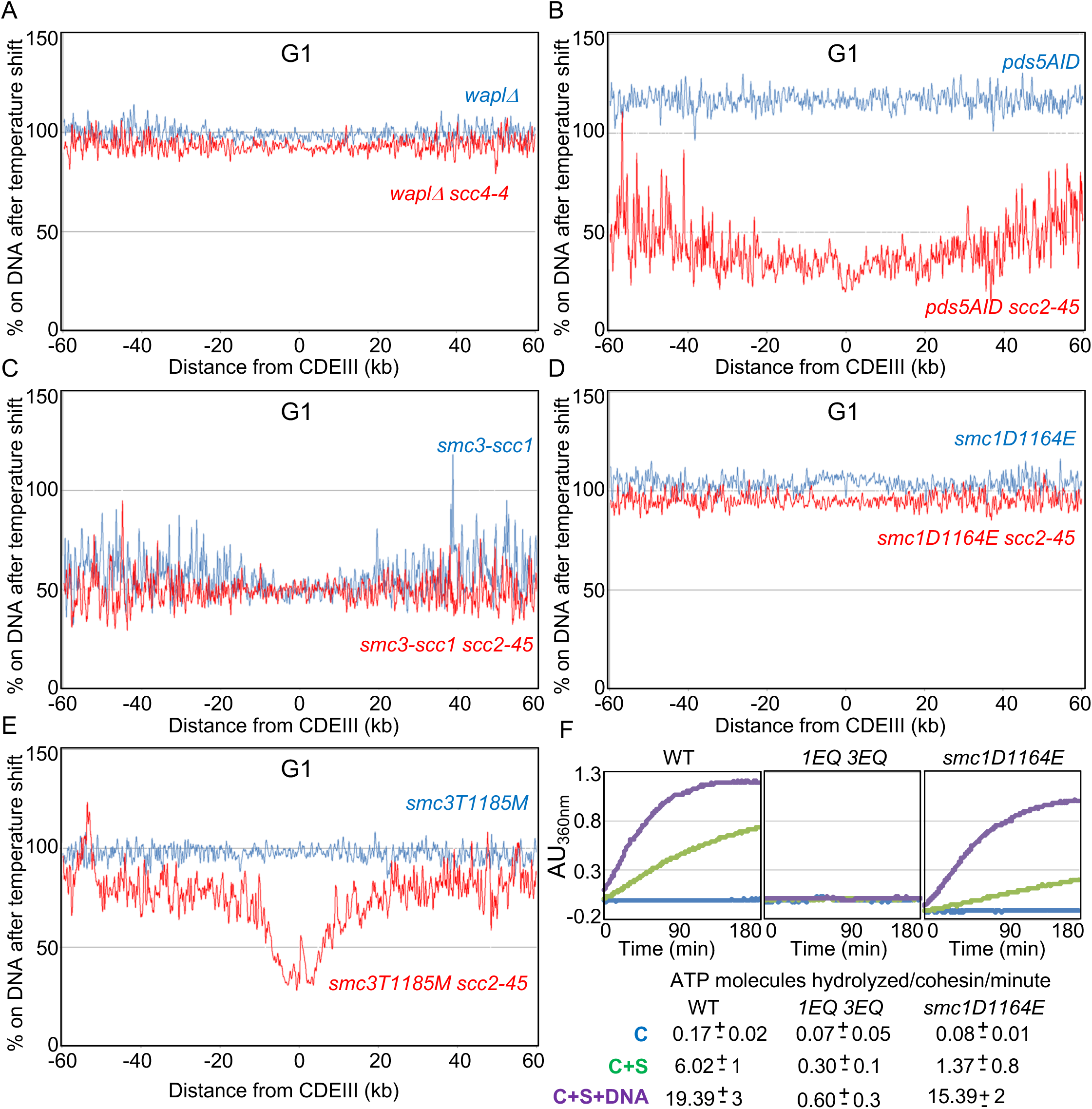
Characterization of the Wapl-independent releasing activity in G1 cells. **(A)** ratio of average cohesin levels before and 20 minutes after the temperature shift of *wpalΔ* (K27569) and *waplΔ scc4-4* (K27570) strains arrested in G1. See Fig S2A. **(B)** *pds5-AID* (K26415) and *pds5-AID scc2-45* (K26414) strains were arrested in G1 with *α*-factor and released into sic1(late G1) arrest in the presence of auxin (IAA) and subjected to temperature shift and Cal-ChIP-Seq. ratio of average cohesin levels before and temperature shift is plotted. **(C)** ratio of average cohesin levels before and 20 minutes after the temperature shift of *smc3-scc1* (K26994) and *smc3-scc1 scc2-45* (K26993) strains arrested in G1. **(D)** ratio of average cohesin levels before and after temperature shift of *smc1D1164E* (K26765) and *smc1D1164E scc2-45* (K26766) strains arrested in G1. **(E)** ratio of average cohesin levels before and after temperature shift of *smc3T1185M* (K27536) and *smc3T1185M scc2-45* (K27537) strains arrested in G1. **(F)** ATPase activity of purified WT, smc1E1155Q smc3E1158Q and smc1D1164E mutant tetramer stimulated by Scc2. The rate of ATP hydrolysis was measured either in the presence or absence of DNA. See Fig S2B.

### Wapl-independent release does not require Pds5

Pds5 is required for Wapl-mediated release and could in principle have a more fundamental role in the release mechanism than Wapl itself. The finding that Scc2 transiently displaces Pds5 from cohesin during the act of loading at centromeres (Petela et al., 2018) suggests that Scc2 might block release by displacing Pds5. If so, depletion of Pds5 from late G1 cells using an auxin-dependent degron should abrogate cohesin’s dissociation from the genome induced by inactivation of Scc2. To test this, we pre-synchronized *PDS5-AID* and *PDS5-AID scc2-45* in early G1 using *α* factor before releasing them in the presence of auxin into a Sic1-mediated late G1 arrest. Calibrated ChIP-seq revealed a major difference in the temperature shift ratio profiles of the two strains (Fig. 2B). Notably, in the absence of Pds5, the temperature shift increased association in *SCC2* (blue) but decreased it in *scc2-45* (red) cells. The significant gap between blue and red curves implies that Scc2 is required to maintain cohesin’s association with chromosomes even in the absence of Pds5. Scc2 cannot therefore block release merely by displacing Pds5.

### Wapl-independent release involves dissociation of the Smc3/Scc1 interface and Smc ATPases

If Wapl-independent release shares a mechanism with Wapl-dependent release, then it should be abrogated by preventing dissociation of the Smc3/Scc1 interface (Chan et al., 2012)(Beckouet et al., 2016). We therefore tested whether cohesin containing an Smc3-Scc1 fusion protein, which restores viability to *eco1Δ* cells by inactivating Wapl-dependent RA, persists on chromosomes upon Scc2 inactivation in late G1 cells. Calibrated ChIP-seq shows that unlike Scc1-PK (Fig. 1E), chromosomal association of an Smc3-Scc1-PK fusion protein is unaffected (Fig. 2C). Therefore, Wapl-independent release blocked by Scc2 in G1 cells involves disengagement of the Smc3/Scc1 interface.

Another property of Wapl-dependent release is its abrogation by mutation of highly conserved residues in Smc1 and Smc3’s ATPases, namely the signature motif *smc1L1129V*, D-loop *smc1D1164E*, and H-loop *smc3T1185M* mutations, which all restore viability to *eco1Δ* cells (Camdere et al., 2015; Elbatsh et al., 2016; Huber et al., 2016). *smc1L1129V* was almost lethal to *scc2-45* cells at 25°C, presumably because its reduced ATPase activity affects loading (Elbatsh et al., 2016), which is already compromised by *scc2-45* (Fig. 1D). In contrast, *smc1D1164E scc2-45* and *smc3T1185M scc2-45* double mutants were sufficiently viable to obtain pure late G1 cultures and test the effect of a temperature shift inactivating *scc2-45*. Calibrated ChIP-seq revealed that inactivation of Scc2 had little effect on chromosomal *smc1D1164E* cohesin (Fig. 2D) and only a modest effect on *smc3T1185M* (Fig. 2E). These data imply that Wapl-independent release revealed by Scc2 inactivation occurs by the same mechanism as release mediated by Wapl in the presence of Scc2.

The abrogation of release by *smc1D1164E* reveals an interesting conundrum. Scc2 activates cohesin’s ATPase and might therefore block release by de-stabilizing the engagement of Smc1/3 ATPase heads. And yet, a release mechanism unleashed by Scc2’s inactivation is eliminated by an *smc1* mutation thought to abrogate release by abolishing ATPase activity (Camdere et al., 2015; Elbatsh et al., 2016). If *smc1D1164E* really eliminated ATP hydrolysis, then Scc2 cannot prevent release by inducing ATP hydrolysis. We therefore purified wildtype (WT), walker B mutants in both Smc1 and Smc3 ATPase (1EQ 3EQ), *smc1D1164E* mutant cohesin tetramers and compared their ATPase activity stimulated by Scc2 in the presence and absence of DNA (Fig. 2F). As expected, the activity of wild type cohesin was fully dependent on Scc2 and stimulated by DNA while the Walker B mutant complex was inactive under all conditions. *smc1D1164E* reduced activity about fourfold, an effect that was largely suppressed by the presence of DNA (Fig. 2F). Thus, suppression of RA by *smc1D1164E* is not necessarily due to an adverse effect on cohesin’s ATPase activity. Our finding that *smc1D1164E* reduced the ability of cohesin containing the *smc3E1155Q* walker B mutant to associate with Scc2 at centromeric loading sites (Fig. S2b) implies that *smc1D1164E* also affects a process prior to ATP hydrolysis.

### Neither Smc3 acetylation nor Pds5 are required to turn off Wapl-independent release

How is Wapl-independent release turned off? Our finding that it shares a similar mechanism to the Wapl-dependent process suggests that Smc3 acetylation may be responsible (Beckouet et al., 2016; Rowland et al., 2009). To test this, we arrested *ECO1 scc3K404E scc2-45* and *eco1Δ scc3K404E scc2-45* cells in G2/M at 25°C and then shifted both to 37°C for 20min. Surprisingly, inactivation of Scc2 had no effect on cohesin’s association with the genome in either culture (Fig. 3A). This was confirmed using sedimentation velocity/gel electrophoresis to measure sister mini-chromosome cohesion, which was unaffected by Scc2 inactivation both in the presence and absence of Eco1 (Fig. 3B). The transition from a state in which Scc2 is required to block Wapl-independent release (G1) to one where it is not (G2) does not therefore require cohesin’s acetylation by Eco1.

**Figure 3.**
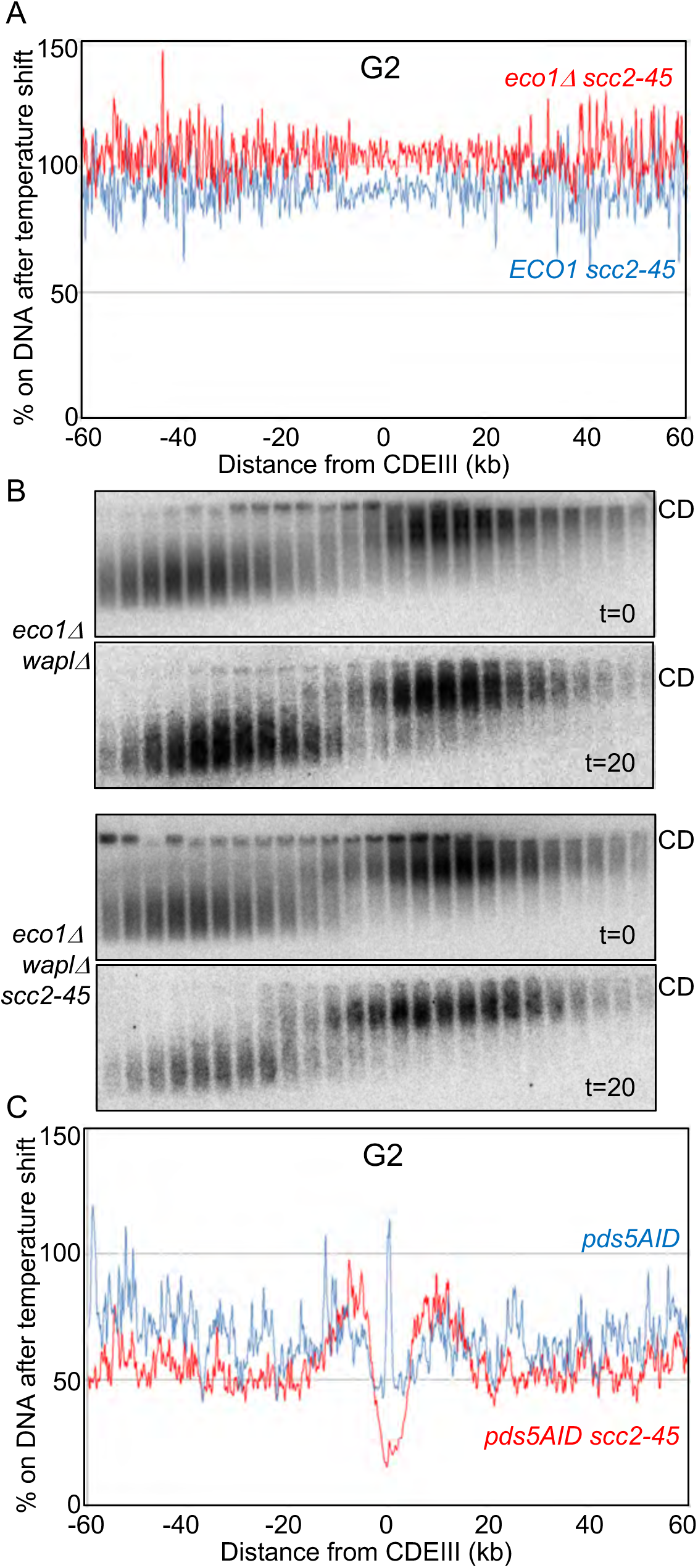
Neither Smc3 acetylation nor Pds5 are required to turn off Wapl-independent release. **(A)** ratio of average cohesin levels before and after temperature shift of *scc3K404E scc2-45* (K24709) and *smc3K404 eco1Δ scc2-45* (K25947) strains arrested in G2. **(B)** *waplΔ eco1Δ* (K22698) and *waplΔ eco1Δ scc2-45* (K22697) strains carrying 2.3kb circular mini-chromosome were arrested in G2 and subjected to temperature shift. Mini-chromosome dimers and monomers were separated by sucrose gradient sedimentation and gel electrophoresis and detected by Southern blotting as detailed in STAR methods. **(C)** *pds5-AID* (K26415) and *pds5-AID scc2-45* (K26414) strains were arrested in G1 with *α*-factor and released into G2 arrest in the presence of auxin (IAA) and subjected to temperature shift and Cal-ChIP-Seq. ratio of average cohesin levels before and temperature shift is plotted.

Because Pds5 is necessary to maintain cohesion, even in the absence of Wapl, it is conceivable that changes in Pds5’s behaviour might be involved. To test this, we synchronised *PDS5-AID* or *PDS5-AID scc2-45* cells in G1 using *α* factor and then allowed them to undergo replication in the presence of IAA and nocodazole. Calibrated ChIP-seq showed that shifting cells to 37°C for 20 min. modestly reduced cohesin’s chromosomal association (Fig. 3C) but importantly, there was little difference between *SCC2* (blue) and *scc2-45* (red) cells. Thus, Wapl-independent release is active in late G1 but not in G2/M cells lacking Pds5 (compare Figs. 2B & 3C). Note that Pds5 is required for Smc3 acetylation and the fact that Pds5 depletion does not abolish Wapl-independent release confirms that acetylation is unnecessary for this process.

### Neither cohesion establishment nor passage through S phase is required to turn off Wapl-independent release

To address whether establishment of cohesion or passage through S phase is required for the switch, we asked whether Scc2 is required to prevent release of cohesin loaded onto chromosomes during G2. *scc3K404E* and *scc3K404E scc2-45* cells containing an ectopic copy of a PK tagged *SCC1* gene under control of the *GAL* promoter were arrested in G2 at 25°C and a pulse of Scc1-PK produced by transient induction with galactose for 60 min. Scc1-PK synthesis was subsequently blocked by transferring cells to glucose, after which the temperature was shifted to 37°C for 20 min. Due to cysteines within *SMC1, SMC3*, and *SCC1-PK* alleles, the Scc1-PK tagged cohesin produced by this protocol was 6C, which permitted measurement of mini-chromosome CMs and CDs. This revealed that cohesin which had loaded during G2 in the absence of conventional releasing activity (due to *scc3K404E*) formed CMs but few if any CDs. As expected, formation of CMs during G2 depended on Scc2 (Fig. S3A). Crucially, these CMs were largely unaffected by Scc2’s inactivation in *scc2-45* cells (Fig. 4A), implying that cohesin loaded onto chromosomes during G2 without forming cohesion does not depend on Scc2 for its maintenance and is therefore not subject to Wapl-independent release. Calibrated ChIP-seq revealed that cohesin along chromosome arms was similarly unaffected by Scc2’s inactivation while peri-centric cohesin was only modestly reduced (Fig. 4B). Thus, cohesin’s switch to a stable form (resistant to Scc2 inactivation) does not require prior association with chromosomes during S phase or indeed establishment of cohesion.

**Figure 4.**
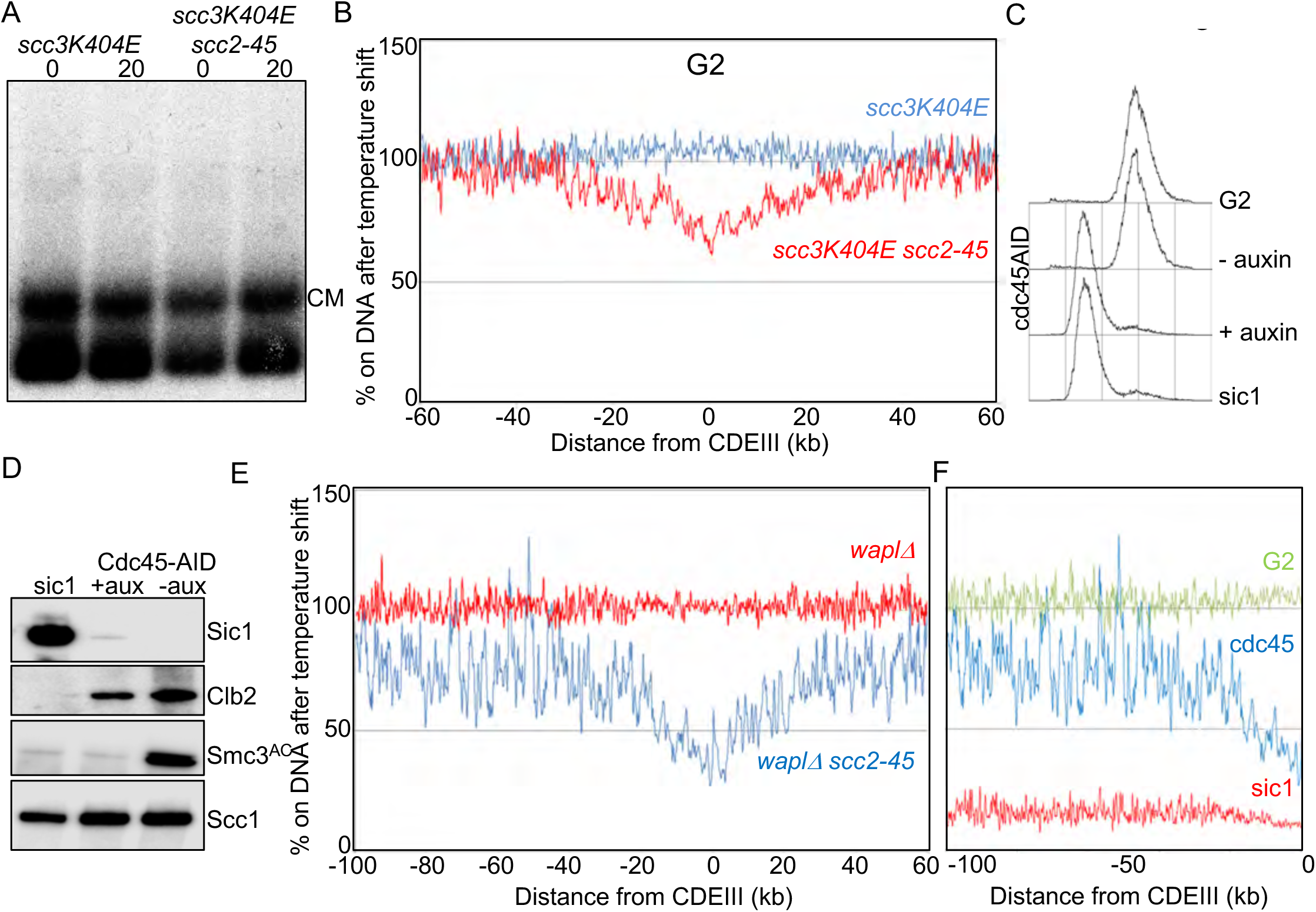
Neither cohesion establishment nor passage through S phase is required to turn off Wapl-independent release. **(A)** *scc3K404E* (K24697) and *scc3K404E scc2-45* (K24738) strains containing 2C Smc1 and 2C Smc3 and galactose inducible 2C Scc1^NC^ were arrested in G2 in YEP raffinose and Scc1 expression induced by addition of galactose. 60 minutes after galactose addiotion, glucose was added to the cultures and temperature shifterd to 37°C. 0 and 20 minute samples were subjected to mini-chromosomeIP (Scc1-PK6). **(B)** 0 and 20 minute samples from (A) were subjected to Cal-ChIP-Seq. ratio of average cohesin levels before and temperature shift is plotted. **(C)** FACS profile of *cdc45-AID* strain (K27169) that was released from a G1 arrest either in the presence or absence of auxin (NAA) and nocodazole is shown along with those of lateG1 and G2 arrested cultures. **(D)** The *cdc45-AID* strain (K27169) grown as in (C) was analysed by western blotting using the indicated antibodies. **(E)** Cdc45 was depleted from *wapl–AID cdc45-AID* (K27169) and *wapl-AID cdc45-AID scc2-45* (K27168) strains like described in (C). Following temperature shift, 0 and 20 minute samples were subjected to Cal-ChIP-Seq. ratio of average cohesin levels before and temperature shift from −100kb to +60KB relative to all 16 centromeres is plotted. **(F)** Re-plotting of ratio of average cohesin levels upon temperature shift resulting in scc2 inactivation upon late G1 arrest (Fig. 1E), G2 arrest (Fig. 1F) and Cdc45 depletion (Fig. 4E). Changes in cohesin levels from −100kb relative to all 16 centromeres is shown.

### DNA replication is not required to switch off Wapl-independent release along chromosome arms

To address whether DNA replication is required for the switch, we analysed the consequences of depleting Cdc45, an essential component of the CMG helicase. *Wapl-AID CDC45-AID* and *wapl– AID scc2-45 CDC45-AID* cells were released from an *α* factor induced G1 arrest in the presence of synthetic auxin (NAA) and nocadozole. Though Cdc45-depleted cells fail to replicate DNA (Fig. 4C) or acetylate Smc3 (Fig. 4D), they nevertheless degrade Sic1, and accumulate modest levels of the mitotic Clb2 cyclin, albeit less than seen in cells allowed to replicate (Fig. 4D). Calibrated ChIP-seq revealed that Scc2 inactivation caused a twofold drop in peri-centric chromosomal cohesin but a more modest change in chromosome arm cohesin (Fig. 4E). Importantly, the effect of inactivating Scc2 in Cdc45-depleted cells more closely resembles the G2 than the G1 pattern (Fig. 4F), at least along chromosome arms, which represents the majority of chromosomal cohesin. These data are consistent with the notion that it is activation of Cdk1 rather than replication per se that switches off cohesin’s release from chromosomes upon Scc2 inactivation.

### Cdk1 is not required for cohesin to persist on chromosomes without Scc2

If Cdk1 is responsible for switching off release when cells initiate S phase, then its inhibition should cause G2 CMs to revert to a state that requires Scc2 for their maintenance. To test this, we used the analogue sensitive *cdc28-as1* allele that can be inhibited in a highly specific manner by addition of the ATP analogue 1NMPP1 (Bishop et al., 2000). *cdc28-as1* cells were arrested in G2/M with nocadazole whereupon Cdk1 was inhibited by addition of 1NMPP1. As expected, this caused Clb2 degradation and Sic1 accumulation, which ensures the complete inhibition of Clb/Cdk1 kinases (Fig. S3B). However, as previously observed (Amon, 1997), these events were accompanied by rapid degradation of Scc1 (Fig.S3B), presumably due to separase activation. To prevent this, we used the *GAL* promoter to express Scc1^NC^-PK, a PK-tagged allele (*scc1R180D R268D*) that cannot be cleaved by separase (Uhlmann et al., 1999). Importantly, the cohesin generated by Scc1^NC^-PK was 6C. *scc3K404E cdc28-as1* and *scc3K404E scc2-45 cdc28-as1* cells were arrested in G1 with *α* factor and then allowed to embark on the cell cycle in the presence of galactose to induce Scc1^NC^-PK and nocadazole to arrest cells in G2. Following completion of S phase, further Scc1^NC^-PK synthesis was shut off, Cdk1 was inhibited by addition of 1NMPP1, and 60 min. later cells were shifted from 25 to 37°C. Remarkably, neither CMs nor CDs produced prior to Cdk1’s inhibition were affected by Scc2’s inactivation (Fig. 5A). The same was also true for CMs made during G2; that is, when Scc1^NC^-PK was induced only after the cells had replicated (Fig. 5B). Thus, Cdk1 activity is not required to maintain chromosomal cohesin in the state resistant to Scc2 inhibition.

**Figure 5.**
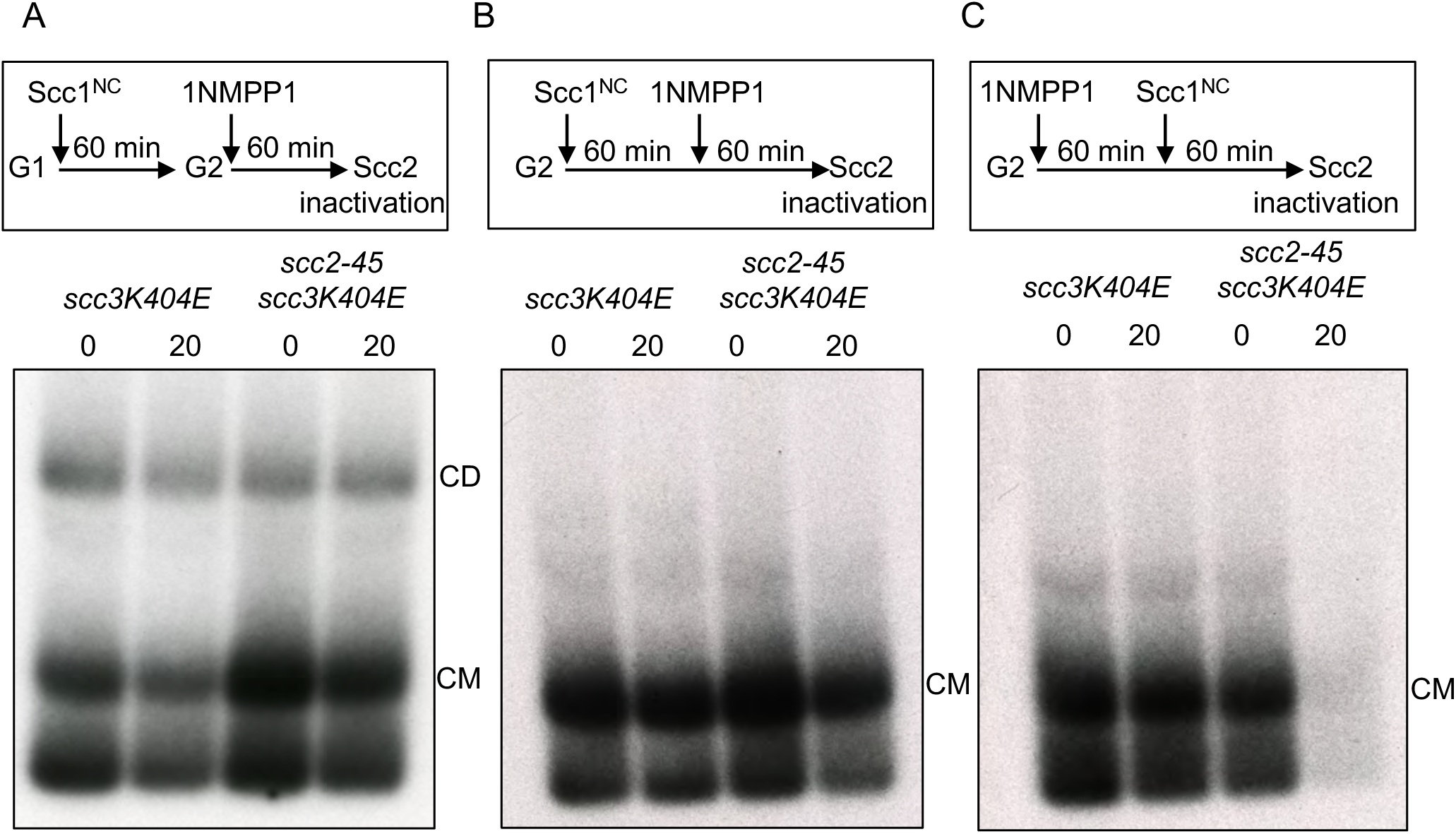
Cdk1 is not required for cohesin to persist on chromosomes without Scc2. **(A)** *scc3K404E cdc28-as1* (K25437) and *scc3K404E scc2-45 cdc28-as1* (K25440) containing galactose inducible 2C Scc1^NC^were released from a G1 arrest into a G2 arrest in the presence of galactose. Glucose and 1NMPP1 for 60 minutes followed by temperature shift and analysis by mini-chromsomeIP. **(B)** Strains described in (A) were arrested in G2 and Scc1^NC^ expression induced by galactose addition for 60 minutes. Glucose and 1NMPP1 were added to the cultures for 60 minutes followed by temperature shift and analysis by mini-chromsomeIP. **(C)** Strains described in (A) were arrested in G2. Followed by 1NMPP1 addition for 60 minutes galactose was added to the cultures to induce 2C Scc1^NC^ for 60 minutes. Glucose was added to the cultures followed by temperature shift and analysis by mini-chromsomeIP. See Fig. S3 for supporting data.

The previous experiment shows that cohesin loaded onto chromosomes in G2 is refractory to Scc2 inactivation even when Clb/Cdk1 is inhibited and yet cohesin loaded onto chromosomes in cells arrested in late G1 with inactive Clb/Cdks is not. If Cdk1 activity were really the parameter that determines whether cohesin requires the Scc2 maintenance function, then the cohesin in these two populations should have behaved identically. One possible explanation is that the low Cdk1 state created by inhibition of Cdk1 in G2 cells is in some way different from the low Cdk1 state created by expression of non-degradable Sic1. In other words, these two cell cycle states in fact differ in some unknown way that affects cohesin’s behaviour. If this is the case, even cohesin loaded onto chromosomes after Cdk1 had been inhibited in G2 cells would behaver differently to that loaded in Sic1-arrested cells.

To address this, we arrested *scc3K404E SCC2 cdc28-as1* and *scc3K404E scc2-45 cdc28-as1* cells in G2, inhibited Cdk1 by 1NMPP1, and then only subsequently induced Scc1^NC^-PK to generate CMs. Crucially, these CMs, unlike those made prior to Cdk1 inhibition (Fig. 5B), disappeared in the *scc2-45* but not in the *SCC2* cells upon shift from 25 to 37°C (Fig. 5C). This implies that the G1-like state created by inhibition of Cdk1 in G2 cells is in fact similar to that produced by arresting cells in late G1 with non-degradable Sic1. CMs made in both states require Scc2 for their maintenance.

This suggests an alternative explanation for the persistence of a G2 character following Cdk1 inhibition of CMs made in G2 cells, namely that their resistance to Scc2 inhibition is conferred by their continued association with chromosomes, and not by cell cycle state per se. In other words, loss of Scc2 resistance characteristic of G2 chromosomal cohesin requires its removal from chromosomes. If the resistance is conferred by a post-translational modification promoted by Cdk1, then the modification must persist even when Cdk1 is subsequently inhibited, as long as cohesin remains associated with chromosomes. Though this might seem unlikely, there is in fact a clear precedent for such behaviour, namely the dependence of Smc3 deacetylation by Hos1 on Scc1 cleavage and not the decline in Cdk1 activity that normally accompanies cleavage during anaphase (Beckouet et al., 2010).

### Scc2 is not required during S phase to establish sister chromatid cohesion

The finding that CMs produced during G2 remain refractory to Scc2 inactivation even when Cdk1 is inhibited provided a means of testing whether Scc2 is required to convert these CMs to CDs when a new round of new replication is triggered by Cdk1 reactivation. We therefore arrested *SCC2* or *scc2-45 scc3K404E scc2-45 cdc28-as1* cells in G2, induced Scc1^NC^-PK to create CMs (Fig. 6D) and subsequently inhibited Cdk1 for 60 min to induce formation of pre-replication complexes (Pre-RCs) (Dahmann et al., 1995). At this point, both strains were shifted to 37°C for 20 min, filtered, and aliquots inoculated into 37°C nocodazole medium with and without 1NMPP1 (Fig. 6A). Remarkably, the DNA re-replication induced by Cdk1 re-activation upon removal of 1NMPP1 (Fig. 6B and C) was accompanied by appearance of CDs in *scc2-45* as well as *SCC2* cells (Fig. 6E).

**Figure 6.**
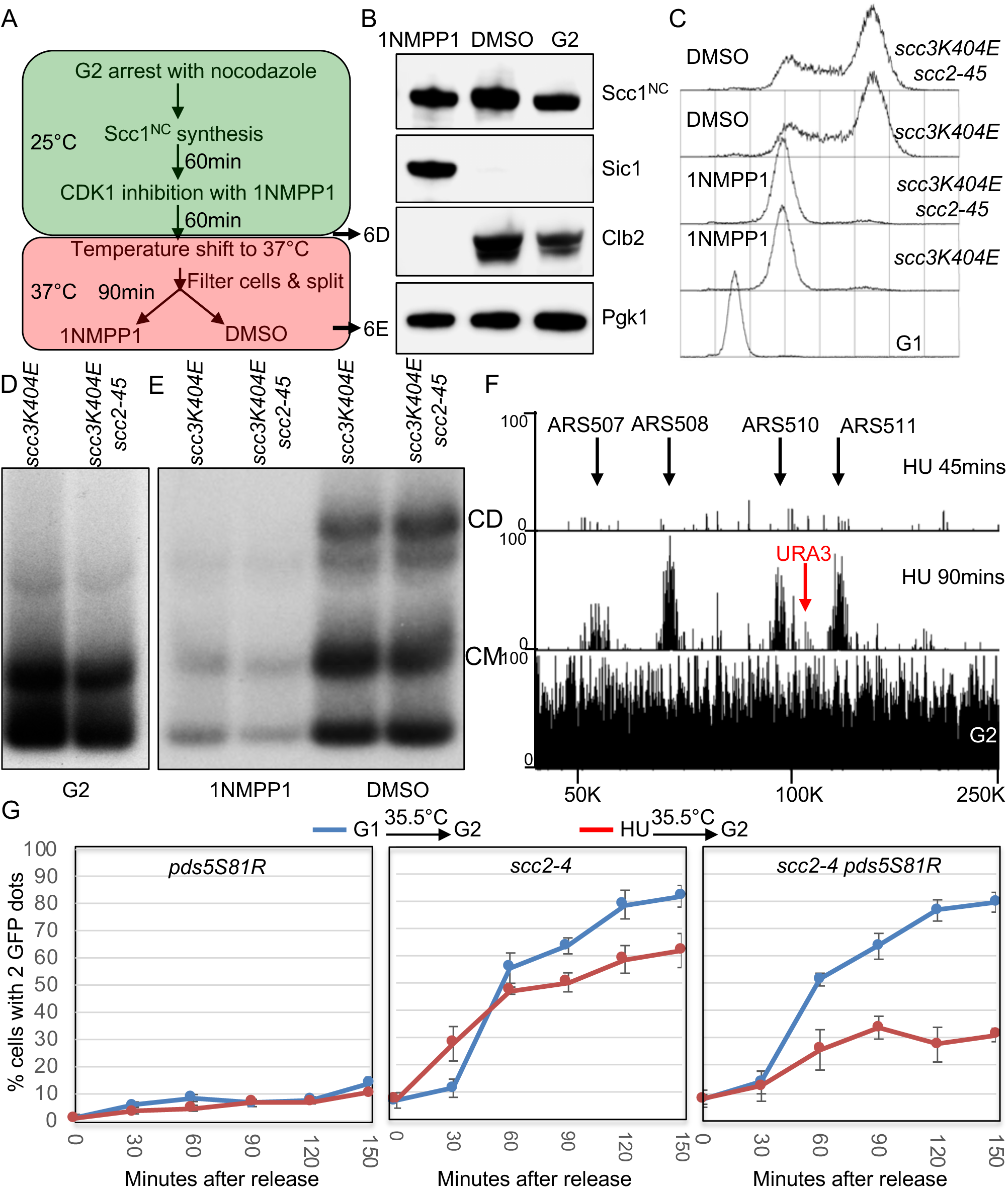
Scc2 is not required during S phase to establish sister chromatid cohesion. **(A)** Schematic summary of the growth conditions for the experiments described below (B-E). **(B)** *scc3K404E cdc28-as1* (K25437) was grown as summarised in (A) and analysed by western blotting with the indicated antibodies. **(C)** FACS profiles of *scc3K404E cdc28-as1* (K25437) and *scc3K404E scc2-45 cdc28-as1* (K25440) strains grown as illustrated in (A). **(D and E)** Mini-chromosomeIP analysis of *scc3K404E cdc28-as1* (K25437) and *scc3K404E scc2-45 cdc28-as1* (K25440) strains grown as illustrated in (A). Samples for (D) and (E) were taken at indicated timepoints in (A). See Fig S4 for supporting data. **(F)** Wild type strain (K699) was released from a G1 arrest into nocadazole or HU containing medium for indicated times and analysed by calibrated genome sequencing as described in STAR methods. A portion of chromosome V is shown with the positions of ARS elements and URA3 gene marked. **(G)** Sister chromatid cohesion was measured in *pds5S81R* (K27443) *scc2-4* (K15028) and *scc2-4 pds5S81R* (K27575) strains that were arrested in G1 or S (HU) and released into G2 arrest (by Cdc20 depletion) at non-permissive temperature.

Because *scc2-45* cells are incapable of forming either CMs or CDs at 37°C when allowed to replicate after release from an *α* factor-induced G1 arrest (Fig. S4A) (Srinivasan et al., 2018), the CDs that appear in *scc2-45* cells when re-replication is induced by transient Cdk1 inhibition (Fig. 6E) were presumably derived from the Scc2-resistant CMs produced during the previous G2 arrest. In other words, CMs can be converted during S phase to CDs in the absence of Scc2 activity. Importantly, the appearance of CDs depended on removal of 1NMPP1 (Fig. 6E), implying that their creation only occurs when re-activation of Cdk1 triggers DNA replication from pre-RCs made during the preceding period of Cdk1 inhibition. Though unexpected, the result was highly reproducible (see Fig. S4B). It is important to note that CMs were not quantitatively converted to CDs in the *scc2-45* cells (Fig. 6E), possibly because of incomplete re-replication (Fig. 6C) (Fig. S4C).

Because of the unexpected results of this experiment and because it merely addressed CD formation by small circular mini-chromosomes, we sought an alternative way of addressing Scc2’s role during S phase. Since Scc2 is required to maintain cohesin on chromosomes in cells that have not yet activated Clb/Cdk1, we treated the cells with hydroxyurea (HU) to delay DNA replication under conditions in which cells activate Cdk1 and therefore no longer require Scc2 to maintain chromosomal cohesin.

To measure cohesion on a chromosome arm, we used a version of chromosome V in which multiple tandem TetO arrays at the *URA3* locus are marked by TetR-GFP (Michaelis et al., 1997). *SCC2* or *scc2-4* cells were released from an *α* factor induced G1 arrest into HU containing medium at 25°C. After 45 min, cells were transferred to HU-free medium at 35.5°C under conditions in which cells were depleted for Cdc20, which prevented separase activation and caused metaphase arrest. Crucially, calibrated whole genome sequencing revealed little or no origin firing during the 45 min incubation in the presence of HU (Fig. 6F).

To assess the ability of these cells (Fig. 6G red graphs) to build cohesion, they were compared to a different population of cells that were released from *α* factor into HU-free medium directly at 35.5°C (Fig. 6G blue graphs). In *SCC2* cells, the fraction of cells with two GFP dots (a measure of cohesion loss) remained low throughout the time course whether or not they had been given an opportunity to load cohesin in the presence of HU at 25°C (data not shown but see *SCC2 pds5S81R* cells in left panel Fig. 6G). Thus, as expected, wild type cells established cohesion under both regimes.

In *scc2-4* cells, the fraction of cells with two GFP dots rose soon after replication, indicating a failure to establish sister chromatid cohesion. Crucially, the cohesion defect was only modestly reduced when cells were allowed to load cohesin onto chromosomes in the presence of HU at 25°C (compare blue and red curves in middle panel Fig. 6G). This implies that loading of cohesin onto chromosomes during the HU arrest at 25°C is insufficient to create efficient cohesion when cells are released from the HU arrest at 35.5°C. Due to Cdk1 activation, Wapl-independent release should be inactive in HU arrested cells but Wapl-dependent release should still take place as very little Smc3 acetylation occurs during the 45 min HU arrest (Nasmyth, 2017).

To test whether Wapl-dependent release contributes to the lack of cohesion establishment, we introduced the *pds5S81R* mutation, which compromises Wapl’s ability to bind Pds5 (Rowland et al., 2009). Remarkably, transient incubation in HU at 25°C in *scc2-4 pdss5S81R* cells greatly ameliorated their cohesion defects following replication at 35.5°C (compare red and blue curves in Fig. 6G right panel). Cohesion was not however restored to the level in *scc2-4 pds5S81R* cells grown only at permissive temperature (Fig. S4D). We conclude that when both Wapl-dependent and independent release mechanisms are inactivated, chromosomal cohesin associated with unreplicated genomes can generate substantial sister chromatid cohesion during S phase in the absence of any further Scc2 activity.

## Discussion

The work described here requires a major re-appraisal of the roles of Wapl and Scc2 in determining the dynamics of cohesin’s association with chromosomes in yeast. First and foremost, Scc2 is not merely a factor that loads cohesin onto chromosomes. In G1 cells lacking Wapl, where chromosomal cohesin does not in fact turnover (Chan et al., 2012), release is nevertheless possible but actively suppressed by Scc2. Because it occurs in the absence of Wapl, we refer to cohesin’s dissociation upon Scc2 inactivation during G1 as Wapl-independent release. Because both types are blocked by fusing Smc3 to Scc1 and abrogated by *smc1D1164E*, we presume that Wapl-dependent and -independent pathways share a molecular mechanism, which involves transient dissociation of Scc1’s NTD from Smc3’s coiled coil. Previous work on mammalian cells showed that Scc2/Nipbl associates transiently but continuously with chromosomal cohesin (Rhodes et al., 2017a). Our work suggests that this association is functionally important.

Our second insight is that contrary to prevailing opinion, neither Wapl nor indeed Pds5 are intrinsic features of cohesin release. In G1 cells, Wapl merely facilitates an activity associated with the ATPases of tripartite rings that is inhibited by Scc2. Our finding that Wapl-independent release is abrogated by *smc1D1164E* suggests that Scc2 functions by altering the behaviour of Smc1/3 ATPase heads. In other words, *SCC2* and *smcD1164E* might abrogate release through a common mechanism. One problem in this regard is the suggestion that *smc1D1164E* abolishes release by abolishing cohesin’s ATPase activity (Camdere et al., 2015; Elbatsh et al., 2016). If correct and Scc2 did likewise, then one would have to postulate that Scc2 acts as an inhibitor as well as an activator of cohesin’s ATPase (Petela et al., 2018), which is difficult to envisage. Our finding that *smc1D1164E* does not in fact abolish cohesin’s ATPase activity raises the possibility that the mutation in fact abrogates release by some other mechanism, which avoids the above conundrum.

We suggest that release involves a form of ATPase head engagement that lasts sufficiently long to create a DNA exit gate (Fig. 7A) and that *smc1D1164E* is defective in attaining or maintaining this state. If so, Scc2 could prevent release merely by stimulating ATP hydrolysis, which would disrupt this “release competent” engaged state before the exit gate actually opens (Fig. 7B). An alternative is that Scc2 blocks release by entrapping DNAs between engaged heads and hinges, which would prevent their escape when the exit gate is opened (Fig. 7C). A final possibility is that Scc2 blocks the exit gate opening that occurs upon head engagement in Scc2’s absence (Fig. 7D).

**Figure 7.**
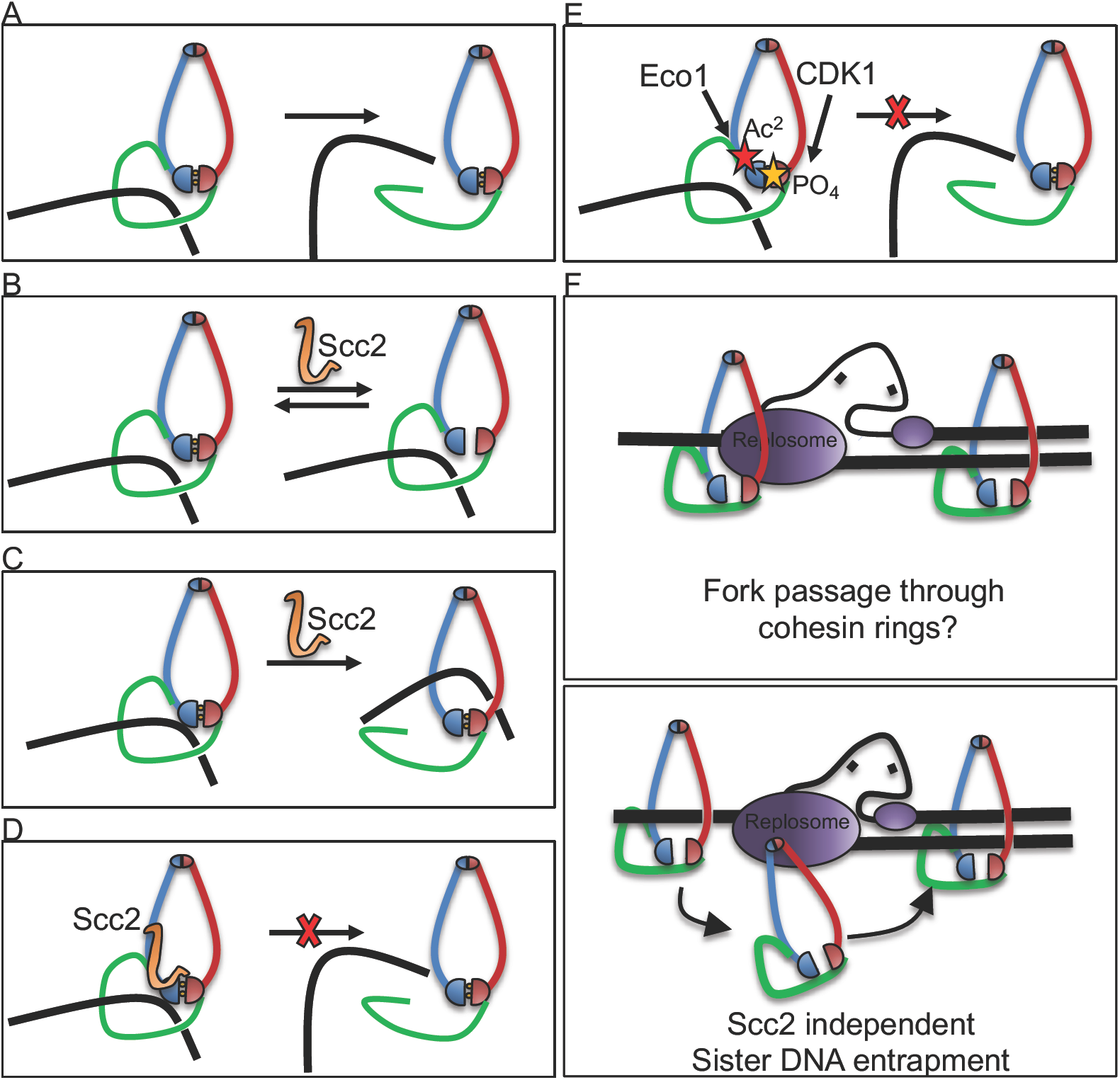
Summary. **(A-D) Scc2 is required in G1 to counteract a Wapl independent releasing activity.** Possible mechanisms of suppression of cohesin release by Scc2. **(E) Increase in CDK1 levels abrogates the Wapl independent release**. This makes Scc2 dispensable for maintaining cohesin’s stable DNA association in G2. **(F) Models for establishment of sister chromatid cohesion.** Scc2 is dispensable for cohesin’s ability to co-entrap sister DNAs during replication. See Discussion for details.

Whether Wapl promotes release in wild type G1 cells by counteracting Scc2’s inhibition of release is unclear. In G2 *eco1 wpl1* cells, Wapl-independent release (as defined above) no longer takes place and yet Wapl induction causes cohesin’s rapid release in these cells. Thus, during G2 at least, Wapl can trigger release by a mechanism that cannot involve counteracting Scc2. Either Wapl triggers release during G1 and G2 by slightly different mechanisms or Wapl does not in fact trigger release during G1 by counteracting Scc2, even though it has the effect of doing so.

The third insight is the discovery that cohesin’s dynamics are subject to a novel type of cell cycle control. Wapl-independent release is specific to G1 cells and is inactivated as cells undergo S phase, not by Smc3 acetylation, but by Clb/Cdk1 activity. Because this regulation does not involve Pds5, Scc2, or Wapl, we suggest that it is mediated by phosphorylation either of Smc-kleisin rings or Scc3. We propose that phosphorylation by Cdk1 as well as acetylation by Eco1 prevent exit gate opening during G2/M (Fig. 7E). We note that there are similarities between this phenomenon and the recent finding that in the absence of Eso1 (Eco1), Wapl induces loss of cohesion in G2 *S.pombe* cells by a pathway involving de-phosphorylation of Rad21 (Scc1) (Birot et al., 2017). Thus, in both organisms cohesin phosphorylation may be accompanied by a change in its dynamics. Though similar in this regard, a key difference is that the release regulated by phosphorylation is Wapl-independent in *S.cerevisiae* but Wapl-dependent in *S.pombe*.

The fourth insight stems from the very different behaviour of newly synthesised and pre-existing chromosomal cohesin in G2 cells that have been converted to a G1-like state through Cdk1 inhibition. While the former requires Scc2 to remain on chromosomes the latter does not. In other words, chromosomal cohesin retains its “G2” behaviour even when a cell’s cell cycle regulatory network is switched to a low Cdk1 G1-like state. This persistence is analogous to abnormal retention of Smc3 acetylation caused by a failure to cleave Scc1.

The fifth and last key insight is our discovery that once cohesin has loaded onto chromosomes in the absence of Wapl, Scc2 is no longer necessary for establishing sister chromatid cohesion. In other words, it is not required for entrapment either of leading or lagging strands during the passage of replication forks. We cannot at this stage rule out the possibility that cohesin associated with unreplicated DNA is in fact displaced by advancing forks but immediately re-associates with both sister DNAs. The key point is that if such displacement and re-association does take place, displacement does not result in diffusion away from the fork (Rhodes et al. 2018) while re-association occurs via an Scc2-independent mechanism. The on rate for cohesin loading onto chromosomes de novo is approximately 33 minutes (Hansen et al., 2017). If the kinetics of re-association at forks obeyed the same rules, then it would be impossible for displaced cohesin to re-associate before it had diffused far away. Thus, if displacement at forks does take place, the kinetics of re-association must be very different to those that normally apply to nucleoplasmic cohesin.

There are two types of explanation for our finding (Fig. 7F). Either replication forks pass through cohesin rings or they open them up in a manner that either does not lead to dissociation or that is associated with an extremely rapid Scc2-independent re-association, albeit with both sister DNAs. Crucially, the mechanism of cohesion establishment during S phase differs in fundamental ways from that of cohesin loading. Not only is the former Scc2-independent but also it is uniquely abrogated by specific mutations in cohesin’s Smc1/3 hinge (Kurze et al., 2011; Mishra et al., 2010).

Though our findings do not exclude the possibility that an Scc2-dependent pathway involving entrapment of single stranded DNA associated with lagging strands (Murayama et al., 2018) co-exists with the Scc2-independent one, they are inconsistent with the claim that the former is essential. On the other hand, the finding that cohesin loading in mammalian cells is dependent on Mcm2-7 during S phase but not during telophase (Zheng et al., 2018) may be pertinent to our finding that entrapment of leading or lagging strands during DNA replication differs from the process of DNA entrapment at other stages of the yeast cell cycle.

## Author Contributions

Madhusudhan Srinivasan, Conceptualization, Data curation, Formal analysis, Validation, Investigation, Visualization, Methodology, Writing—original draft, Project administration, Writing—review and editing. Naomi Petela, Johanna C. Scheinost, James Collier, Menelaos Voulgaris, Maurici Brunet-Roig, Frederic Beckouët: Formal analysis, Methodology, Data curation. Bin Hu: Conceptualization, Formal analysis, Funding acquisition. Kim Nasmyth: Conceptualization, Supervision, Funding acquisition, Writing—original draft, Project administration, Writing—review and editing.

## Acknowledgements

Maria Demidova conducted initial experiments that this study expanded on. We are grateful to Tomo Tanaka and Seiji Tanaka for supplying reagents. We thank all members of the Nasmyth group for valuable discussions, technical assistance and critical reading of the manuscript. This work was funded by the Wellcome Trust Senior Investigator Award, Grant Ref 107935/Z/15/Z and Cancer Research UK Programme Grant, Grant Ref 26747 to K.N. B.H is funded by (202062/Z/16/Z)

## Supplementary figure legends

**Figure Supplement 1.**
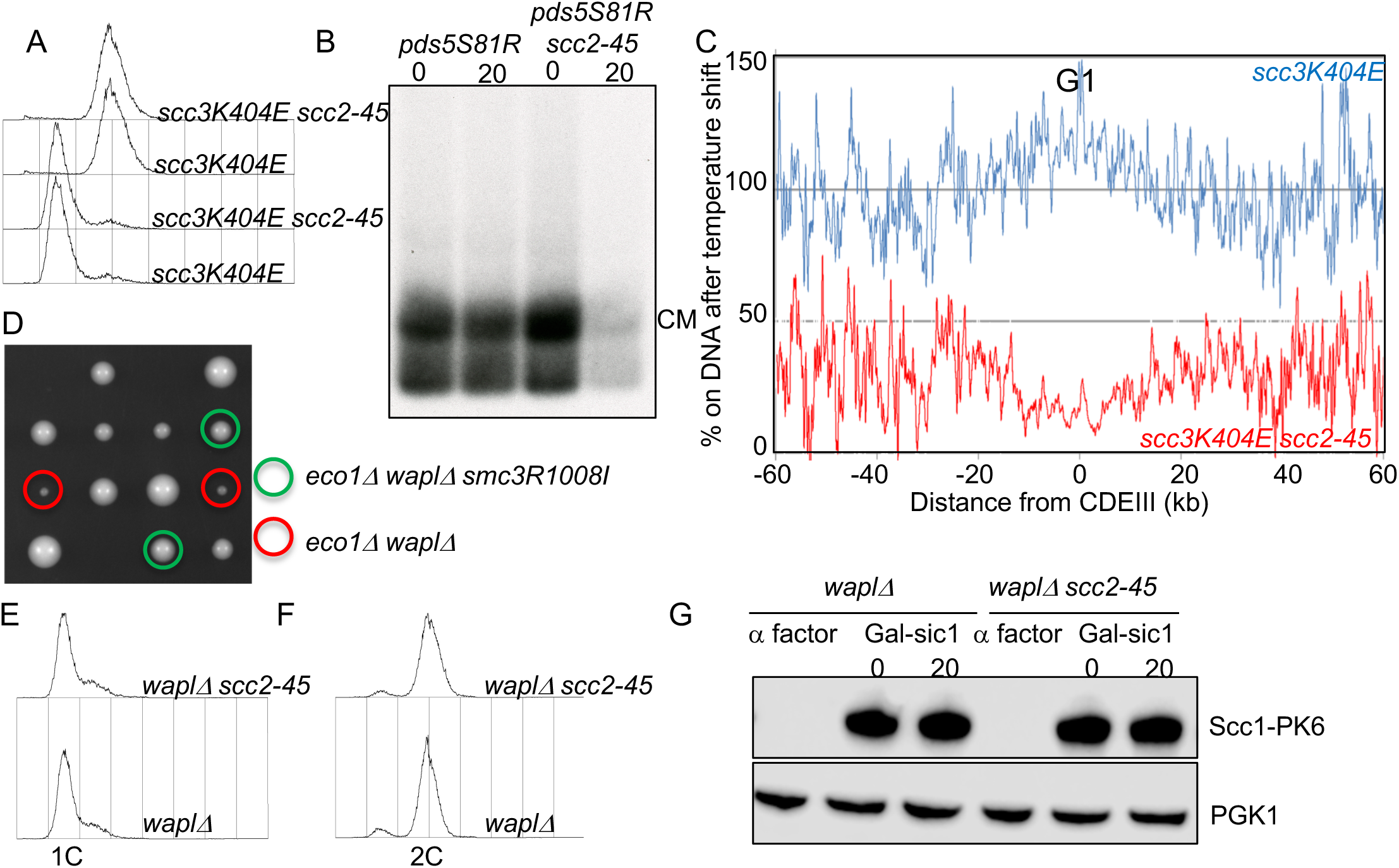
A wapl independent activity releases cohesin from DNA in G1, Supplement to Figure 1. **(A)** FACS profile of the strains used in the experiment described in Figure 1C. **(B)** *pds5S81R* (K25311) and *pds5S81R scc2-45* (K25312) 6C strains expressing galactose-inducible nondegradable sic1 were arrested in late G1 at 25°C as described in STAR methods. The cultures were shifted to 37°C for 20 minutes, aliquots drawn before (0) and after (20) temperature shift were subjected to mini-chromosome IP. **(C)** *scc3K404E* (K25313) and *scc3K404E scc2-45* (K25316) strains used in Figure 1C were arrested in late G1 at 25°C and subjected to temperature shift to 37°C for 20 minutes. 0 and 20 minute samples were analyzed by calibrated ChIP-sequencing (Scc1-PK6). The ratio of average cohesin levels 60 kb on either side of all 16 centromeres before and 20 minutes after the temperature shift is shown. **(D)** Haploid segregants following tetrad dissection of asci from diploid strain (*eco1Δ/ECO1 waplΔ/WAPL smc3R1008I/SMC3*). **(E & F)** FACS profile of the strains used in the experiment described in Figure 1E & F. **(G)** In the experiment described in Figure 1E, samples of the *waplΔ* and *waplΔ scc2-45* were drawn when cells were arrested in G1, late G1 and 20 minutes after the temperature shift in late G1. Cell pellets were lysed and the samples analysed by western blotting against anti PK antibody (Scc1) and anti PGK1 antibody.

**Figure Supplement 2.**
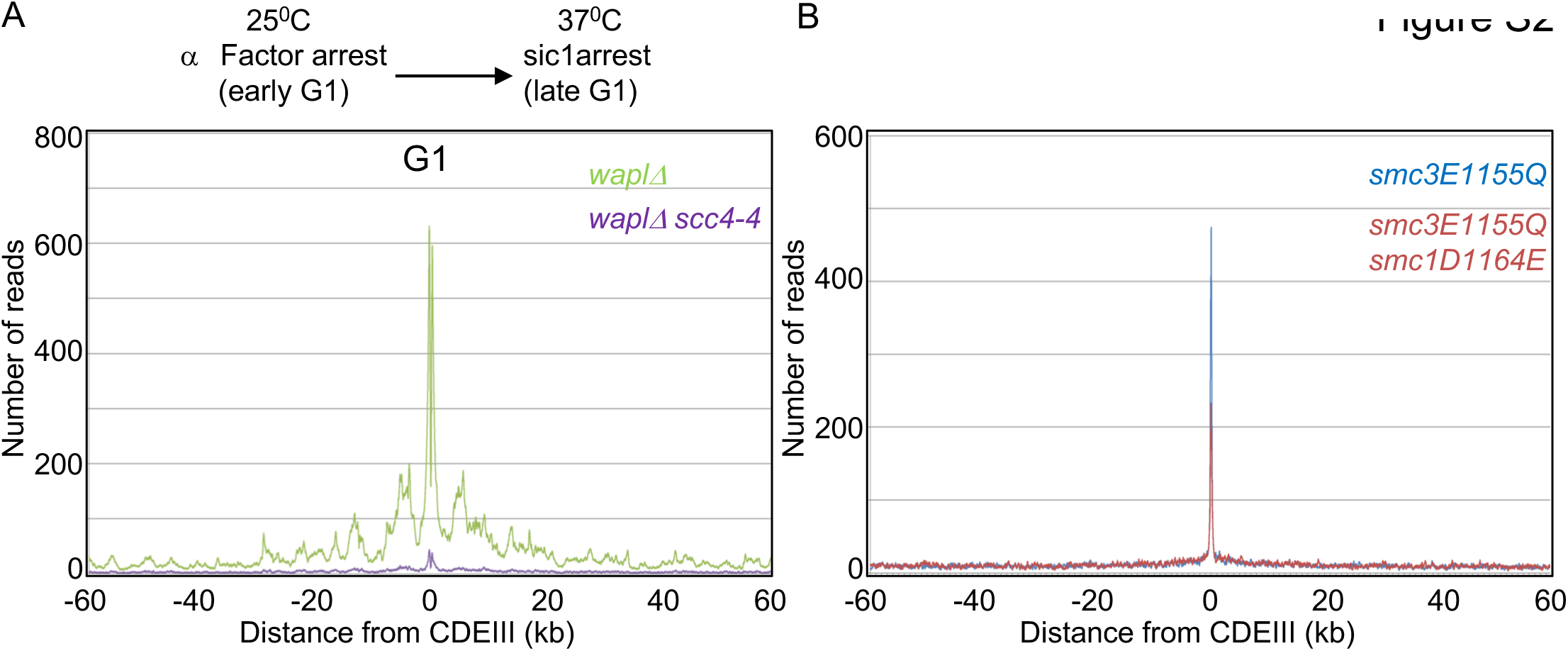
Scc4 is essential for loading cohesin in late G1 and Smc1 D-loop mutant affects cohesin loading, Supplement to Figure 2. **(A)** *wpalΔ* (K27569) and *waplΔ scc4-4* (K27570)described in Figure 2A were arrested in G1 at 25°C and released into late G1 arrest for 90 minutes at 37°C. Samples drawn after 90 minutes in late G1 were analyzed by calibrated ChIP-sequencing (Scc1-PK6). Cohesin ChIP profiles showing the number of reads at each base pair away from the CDEIII element averaged over all 16 chromosomes is shown. **(B)** Calibrated ChIP profile of *smc3E1155Q* (K17409) and *smc3E1155Q smc1D1164E* (25039). Cohesin ChIP profiles showing the number of reads at each base pair away from the CDEIII element averaged over all 16 chromosomes is shown.

**Figure S3.**
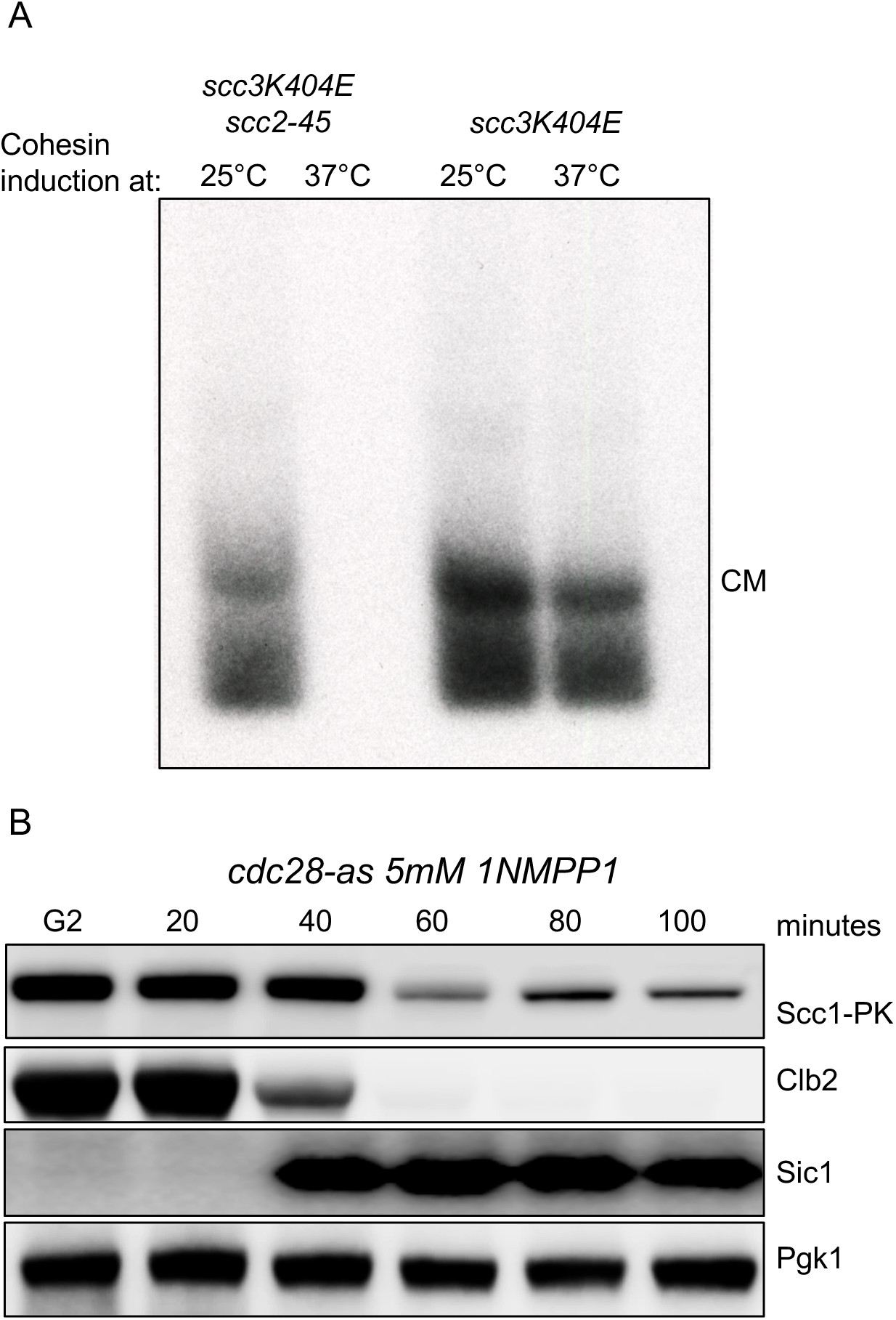
CDK1 inhibition in G2 leads to Scc1 degradation. Supplement to Figures 4 and 5. **(A)** *scc3K404E* (K24697) and *scc3K404E scc2-45* (K24738) strains containing 2C Smc1 and 2C Smc3 and galactose inducible 2C Scc1^NC^ were arrested in G2 in YEP raffinose. The culture was split into two, one half incubated at 25°C and the other at 37°C. Scc1 expression induced by addition of galactose. 60 minutes after galactose addition samples drawn and the pellets were subjected to mini-chromosomeIP (Scc1-PK6). **(B)** *cdc28-as1*(K25423) cells were arrested in G2 and treated with 5μM 1NMPP1, samples were drawn at indicated times and subjected to western blot analysis with the indicated antibodies.

**Figure S4.**
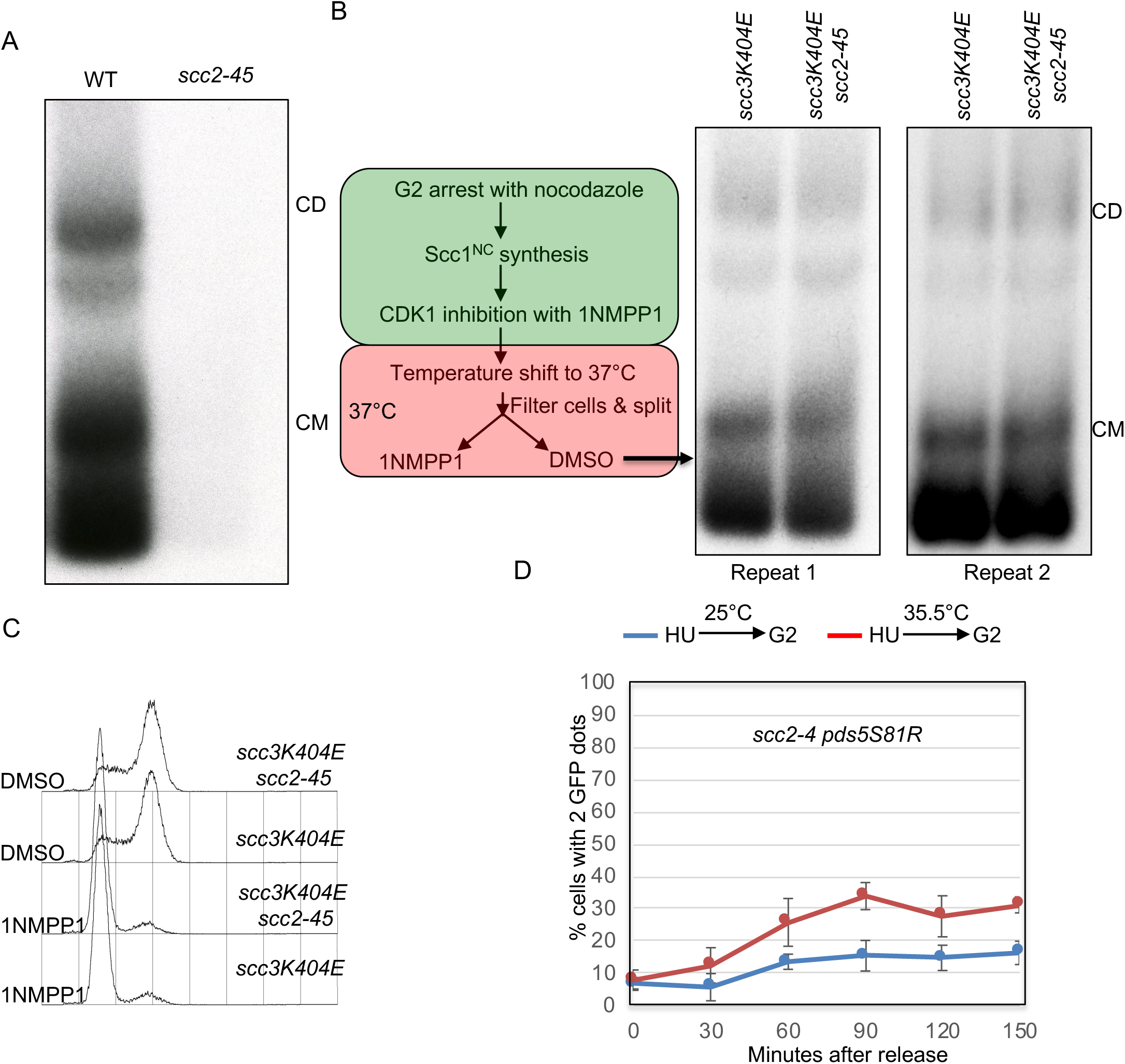
Scc2 independent cohesion establishment. Supplement to Figure 6. CMs and CDs in WT (K23889) and scc2-45 (K24267) 6C strains arrested in G1 with α factor at 25°C in YPD medium and released into nocodazole at 37°C. Two independent repetitions of the experiment described in Figure 6D. **(C)** FACS profiles of *scc3K404E cdc28-as1* (K) and *scc3K404E scc2-45 cdc28-as1* (K) strains grown as illustrated in (B). **(D)** Sister chromatid cohesion was measured in *scc2-4 pds5S81R* strain (K27575) that was arrested in S (HU) and released into G2 arrest (by Cdc20 depletion) at either permissive (blue curve) or non-permissive temperature (red curve).

## Materials and Methods

### CONTACT FOR REAGENT AND RESOURCE SHARING

Further information and requests for resources and reagents should be directed to and will be fulfilled by the lead contact Kim Nasmyth.

### EXPERIMENTAL MODELS

#### Yeast cell culture

All strains are derivatives of W303 (K699). Strain numbers and relevant genotypes of the strains used are listed in the Key Resource Table. Cells were cultured at 25°C in YEP medium with 2% glucose unless stated otherwise. To arrest the cells in G1, α-factor was added to a final concentration of 2 mg/L, every 30 min for 2.5 h. Cells were released from G1 arrest by filtration wherein cells were captured on 1.2 μm filtration paper (Whatman^®^ GE Healthcare), washed with 1 L YEPD and resuspended in the appropriate fresh media. To inactivate Scc2 (*scc2-45*: temperature sensitive allele), fresh media was pre-warmed prior to filtration.

To arrest cells in G2, nocodazole (Sigma) was added to the fresh media to a final concentration of 10 μg/mL and cells were incubated until the synchronization was achieved (>95% large-budded cells).

Cells were arrested in late G1 by galactose-induced overexpression of a non-degradable mutant of the Sic1 protein (mutation of 9 residues phosphorylated by Cdk1). To achieve this, cells were grown in YEP supplemented with 2% raffinose and arrested in G1 as described above. 1 h before release from G1 arrest, galactose was added to 2% of the final concentration. Cells were released into YEPD as described above, and incubated for 60 min at 25°C.

For auxin induced degradation of proteins, cells were arrested in G1 as above and 1 h prior to release auxin (indole-3-acetic acid sodium salt; Sigma) was added to a final concentration of 1 mM. Cells were released from G1 arrest into YEPD medium containing 1 mM auxin and 10 μg/mL nocodazole.

To inhibit CDK1 *cdc28-as1* cells were arrested in G2 with nocodazole (Sigma) until synchronization was achieved (>95% large-budded cells) at 25°C. Subsequently, 1NMPP1 (5μM final) was added and the cultures incubated for 60 minutes at 25°C.

To induce re-replication, Cultures where CDK1 was inhibited (as described above) were filtered and washed with 1l fresh YEPD medium, the cells were resuspended in fresh YEPD medium containing nocodazole and wither 1NMPP1 or DMSO and incubated for further 120-150 minutes.

### METHOD DETAILS

#### In vivo chemical crosslinking

Strains were grown in YEPD at 25°C to OD_600nm_ = 0.5-0.6. 12 OD units were washed in ice-cold PBS and re-suspended in 1 mL ice-cold PBS. The suspensions were then split into 2 x 500 μL and 20.8 μL BMOE (stock: 125 mM in DMSO, 5 mM final) or DMSO was added for 6 min on ice. Cells were washed with 2 x 2 mL ice-cold PBS containing 5 mM DTT, resuspended in 500 μL lysis buffer (25 mM Hepes pH 8.0, 50 mM KCl, 50 mM MgSO_4_, 10 mM trisodium citrate, 25 mM sodium sulfite, 0.25% triton-X, freshly supplemented with Roche Complete Protease Inhibitors (2X) and PMSF (1 mM), lysed in a FastPrep-24 (MP Biomedicals) for 3 x 1 min at 6.5 m/s with 500 μl of acid-washed glass beads (425-600 μm, Sigma) and lysates cleared (5 min, 12 k*g*). Protein concentrations were adjusted after Bradford assay and cohesin immuno-precipitated using anti-PK antibody (AbD Serotec, 1 h, 4°C) and protein G dynabeads (1 h, 4°C, with rotation). Beads were washed with 2 x 1 mL lysis buffer, resuspended in 50 μl 2x sample buffer, incubated at 95°C for 5 min and the supernatant loaded onto a either 3-8% Tris-acetate or 4-12% Bis-Tris gradient gels (Life Technologies).

#### Minichromosome IP

Strains containing a 2.3 kb circular minichromosome harbouring the *TRP1* gene were grown overnight in –TRP medium at 25°C and sub-cultured in YEPD medium for exponential growth (OD_600nm_ = 0.6). 30 OD units were washed in ice-cold PBS and processed for in vivo crosslinking as described above with the following modification: after cohesin immuno-precipitation protein G dynabeads were washed with 2 x 1 ml lysis buffer, resuspended in 30 μl 1% SDS with DNA loading dye, incubated at 65°C for 4 min and the supernatant run on a 0.8% agarose gel containing ethidium bromide (1.4 V/cm, 22h, 4°C). After Southern blotting using alkaline transfer, bands were detected using a 32-P labeled TRP1 probe.

#### SDS Gel electrophoresis and Western blotting

Whole cell lysates were resolved in NuPAGE^TM^ 3-8% or 4-12% gradient gels (ThermoFisher Scientific) and transferred onto PVDF membranes using the Trans-blot Turbo transfer system (BioRad). For visualization, the membrane was incubated with Immobilon Western Chemiluminescent HRP substrate (Millipore) before detection using an ODYSSEY Fc Imaging System (LI-COR).

| Primary Antibody | Origin | Dilution |
| --- | --- | --- |
| V5 Tag | Mouse | 1:1000 |
| Sic1 | Rabbit | 1:1000 |
| Clb2 | Rabbit | 1:1000 |
| Smc3<sup>AC</sup> | Mouse | 1:1000 |
| PGK1 | Mouse | 1:5000 |

#### Calibrated ChIP-sequencing

Cells were grown exponentially to OD_600_ = 0.5 and the required cell cycle stage where necessary. 15 OD_600nm_ units of *S. cerevisiae* cells were then mixed with 5 OD_600nm_ units of *C. glabrata* to a total volume of 45 mL and fixed with 4 mL of fixative (50 mM Tris-HCl, pH 8.0; 100 mM NaCl; 0.5 mM EGTA; 1 mM EDTA; 30 % (v/v) formaldehyde) for 30 min at room temperature (RT) with rotation.

The fixative was quenched with 2 mL of 2.5 M glycine (RT, 5 min with rotation). The cells were then harvested by centrifugation at 3,500 rpm for 3 min and washed with ice-cold PBS. The cells were then resuspended in 300 μL of ChIP lysis buffer (50 mM Hepes-KOH, pH 8.0; 140 mM NaCl; 1 mM EDTA; 1% (v/v) Triton X-100; 0.1% (w/v) sodium deoxycholate; 1 mM PMSF; 2X Complete protease inhibitor cocktail (Roche)) and an equal amount of acid-washed glass beads (425-600 μm, Sigma) added before cells were lysed using a FastPrep^®^-24 benchtop homogeniser (M.P. Biomedicals) at 4°C (3 x 60s at 6.5 m/s or until >90 % of the cells were lysed as confirmed by microscopy).

The soluble fraction was isolated by centrifugation at 2,000 rpm for 3 min then sonicated using a bioruptor (Diagenode) for 30 min in bursts of 30 s on/30 s off at high level in a 4°C water bath to produce sheared chromatin with a size range of 200-1,000 bp. After sonication the samples were centrifuged at 13,200 rpm at 4°C for 20 min and the supernatant was transferred into 700 μL of ChIP lysis buffer. 30 μL of protein G Dynabeads (Invitrogen) were added and the samples were pre-cleared for 1 h at 4 °C. 80 μL of the supernatant was removed (termed ‘whole cell extract [WCE] sample’) and 5 μg of antibody (anti-PK (Bio-Rad) or anti-HA (Roche)) was added to the remaining supernatant which was then incubated overnight at 4°C. 50 μL of protein G Dynabeads were then added and incubated at 4°C for 2 h before washing 2x with ChIP lysis buffer, 3x with high salt ChIP lysis buffer (50 mM Hepes-KOH, pH 8.0; 500 mM NaCl; 1 mM EDTA; 1 % (v/v) Triton X-100; 0.1 % (w/v) sodium deoxycholate; 1 mM PMSF), 2x with ChIP wash buffer (10 mM Tris-HCl, pH 8.0; 0.25 M LiCl; 0.5 % NP-40; 0.5 % sodium deoxycholate; 1 mM EDTA; 1 mM PMSF) and 1x with TE pH7.5. The immunoprecipitated chromatin was then eluted by incubation in 120 μL TES buffer (50 mM Tris-HCl, pH 8.0; 10 mM EDTA; 1 % SDS) for 15 min at 65°C and the collected supernatant termed ‘IP sample’. The WCE samples were mixed with 40 μL of TES3 buffer (50 mM Tris-HCl, pH 8.0; 10 mM EDTA; 3 % SDS) and all samples were de-crosslinked by incubation at 65°C overnight. RNA was degraded by incubation with 2 μL RNase A (10 mg/mL; Roche) for 1 h at 37°C and protein was removed by incubation with 10 μL of proteinase K (18 mg/mL; Roche) for 2 h at 65°C. DNA was purified using ChIP DNA Clean and Concentrator kit (Zymo Research).

#### Extraction of yeast DNA for deep sequencing

Cultures were grown to exponential phase (OD600 = 0.5). 12.5 OD600 units were then collected and diluted to a final volume of 45mL before fixation as described in the protocol for ChIP-seq. The samples were treated as specified in the ChIP-seq protocol up to the completion of the sonication step whereby 80 μL of the samples were carried forward and treated as WCE samples.

#### Preparation of sequencing libraries

Sequencing libraries were prepared using NEBNext^®^ Fast DNA Library Prep Set for Ion Torrent^(tm)^ Kit (New England Biolabs) according to the manufacturer’s instructions. Briefly, 10-100 ng of fragmented DNA was converted to blunt ends by end repair before ligation of the Ion Xpress^(tm)^ Barcode Adaptors. Fragments of 300 bp were then selected using E-Gel^®^ SizeSelect^(tm)^ 2 % agarose gels (Life Technologies) and amplified with 6-8 PCR cycles. The DNA concentration was then determined by qPCR using Ion Torrent DNA standards (Kapa Biosystems) as a reference. 12-16 libraries with different barcodes could then be pooled together to a final concentration of 350 pM and loaded onto the Ion PI^(tm)^ V3 Chip (Life Technologies) using the Ion Chef^(tm)^ (Life Technologies). Sequencing was then completed on the Ion Torrent Proton (Life Technologies), typically producing 6-10 million reads per library with an average read length of 190 bp.

#### Data analysis, alignment and production of BigWigs

Unless otherwise specified, data analysis was performed on the Galaxy platform. Quality of reads was assessed using FastQC (Galaxy tool version 1.0.0) and trimmed as required using ‘trim sequences’ (Galaxy tool version 1.0.0). Generally, this involved removing the first 10 bases and any bases after the 200^th^, but trimming more or fewer bases may be required to ensure the removal of kmers and that the per-base sequence content is equal across the reads. Reads shorter than 50 bp were removed using Filter FASTQ (Galaxy tool version 1.0.0, minimum size: 50, maximum size: 0, minimum quality: 0, maximum quality: 0, maximum number of bases allowed outside of quality range: 0, paired end data: false) and the remaining reads aligned to the necessary genome(s) using Bowtie2 (Galaxy tool version 0.2) with the default (--sensitive) parameters (mate paired: single-end, write unaligned reads to separate file: true, reference genome: SacCer3 or CanGla, specify read group: false, parameter settings: full parameter list, type of alignment: end to end, preset option: sensitive, disallow gaps within *n-*positions of read: 4, trim *n*-bases from 5’ of each read: 0, number of reads to be aligned: 0, strand directions: both, log mapping time: false).

To generate alignments of reads that uniquely align to the *S. cerevisiae* genome, the reads were first aligned to the *C. glabrata* (CBS138, genolevures) genome with the unaligned reads saved as a separate file. These reads that could not be aligned to the *C. glabrata* genome were then aligned to the *S. cerevisiae* (sacCer3, SGD) genome and the resulting BAM file converted to BigWig (Galaxy tool version 0.1.0) for visualisation. Similarly, this process was done with the order of genomes reversed to produce alignments of reads that uniquely align to *C. glabrata*.

#### Visualisation of ChIP-seq profiles

The resulting BigWigs were visualized using the IGB browser. To normalize the data to show quantitative ChIP signal the track was multiplied by the samples’ occupancy ratio (OR) and normalized to 1 million reads using the graph multiply function. In order to calculate the average occupancy at each base pair up to 60 kb around all 16 centromeres, the BAM file that contains reads uniquely aligning to *S. cerevisiae* was separated into files for each chromosome using ‘Filter SAM or BAM’ (Galaxy tool version 1.1.0). A pileup of each chromosome was then obtained using samtools mpileup (Galaxy tool version 0.0.1) (source for reference list: locally cached, reference genome: SacCer3, genotype likelihood computation: false, advanced options: basic). These files were then amended using our own script (chr_position.py) to assign all unrepresented genome positions a value of 0. Each pileup was then filtered using another in-house script (filter.py) to obtain the number of reads at each base pair within up to 60 kb intervals either side of the centromeric CDEIII elements of each chromosome. The number of reads covering each site as one successively moves away from these CDEIII elements could then be averaged across all 16 chromosomes and calibrated by multiplying by the samples’ OR and normalizing to 1 million reads.

#### Cohesin Tetramer and Scc2 purification

All versions of the cohesin complexes purified bear a twin StrepII tag on the Scc1 kleisin. This is the same for the Scc2 construct used in this study except the later bears a single Strep-II tag. Typically 500ml of SF-9 insect cells were grown to ∼3 million/ml and infected with the appropriate baculovirus stock in a 1/100 dilution. Infection was monitored daily and cells harvested when lethality (assayed by the trypan blue test) reached no more than 70%–80%. Cell pellets were then frozen in liquid nitrogen and stored at 80°C. Upon thawing, the pellets were suspended in a final volume of ∼65-70ml with Buffer A (final concentrations of: 25mM HEPES pH 8.0, NaCl 150mM, TCEP-HCl 1mM and Glycerol 10%) and the suspension was immediately supplemented with 2 dissolved tablets of Roche Complete Protease (EDTA-free), 75μg of RNase I and 7μl of DNaseI (Roche, of 10U/μl stock). The cells were then sonicated at 80% amplitude for 5 s/burst/35ml of suspension using a Sonics Vibra-Cell (3mm microtip). In total, 12 bursts were given for every 35ml half of the 70ml suspension (the sonication was always performed in ethanolised ice). A spin at 235,000 x g (45,000rpm on a Ti45 fixed angle rotor) followed for 45 mins following addition of PMSF to 1mM final concentration. The isolated cleared extract was supplemented with 2mM EDTA and was then used to load a 2×5ml StrepTrap HP (Fisher Scientific) column at 1ml/min in an ÄKTA Purifier 100. Wash with Buffer A at 1ml/min to the point of Δ?U_280nm_∼0 and protein elution ensued using Buffer A+20mM desthiobiotin (Fisher Scientific) at 1ml/min. Peak fractions were analyzed using SDS-PAGE and were further purified in a Superose 6 Increase 10/300 (VWR) using Buffer A as running buffer (free of EDTA/PMSF). The resulting peaks were again analyzed using SDS-PAGE and the concentration was determined in Nanodrop using A280. Protein was aliquoted and stocked typically in concentrations ranging from 1 to 3mg/ml.

#### ATPase assay

ATPase activity was measured by using the EnzChek phosphate assay kit (Invitrogen) by following the protocol as provided. Cohesin tetramer (Smc1, Smc3, Scc1 and Scc3; final concentration: 50 nM, final NaCl concentration: 50 mM) was added together with a 40 bp long double stranded DNA (700 nM). The reaction was started with addition of ATP to a final concentration of 1.3 mM (final reaction volume: 150 μl). After completion, a fraction of each reaction was run on SDS-PAGE and the gel stained with coomassie brilliant blue in order to test that the complexes were intact throughout the experiment and that equal amounts were used when testing various mutants and conditions.

### QUANTIFICATION AND STATISTICAL ANALYSIS

#### ATPase assay

ATPase activity was measured by recording absorption at 360 nm every 30 s for 90 min using a PHERAstar FS. Δ?U at 360 nm was translated to P_i_ release using an equation derived by a standard curve of KH_2_PO_4_ (EnzChek kit). Rates were calculated from the slope of the linear phase (first 10 min). At least two independent biological experiments were performed for each experiment, means and standard deviations are reported for every experiment.

### DATA AND SOFTWARE AVAILABILITY

#### Scripts

All scripts written for this analysis method are available to download from https://github.com/naomipetela/nasmythlab-ngs.

*Chr_position.py* takes mpileups for *S. cerevisiae* chromosomes and fills in gaps, with each position in the chromosome added given a read depth of 0.

*Filter60.py* reads the files produced by Chr_position.py and takes the read depth for all positions 60 kb either side of the CDEIII for all chromosomes, produces an average for each position and multiples it by the OR. The OR should be derived from the reads aligned in the appropriate bam files (Hu et al., 2015).

#### Calibrated ChIP-seq data

The calibrated ChIP-seq data (raw and analyzed data) have been deposited on GEO under accession number.

|  |  |  |
| --- | --- | --- |
| *S. cerevisiae* MATa *ade2-1 trp1-1 can1-100 leu2-3,112 his3-11,15 ura3 GAL psi+*<br>*All following strains are based on this background* | This study | **K699** |
| *S. cerevisiae* MATa *scc2-4 SCC1-PK9 TETR-GFP::LEU2 TET Os::URA3 trp1::MET3p-CDC20::TRP1* | This study | **K15028** |
| *S. cerevisiae* MATa *trp1::smc3(E1155Q)-PK6::TRP1* | This study | **K17409** |
| *S. cerevisiae* MATa *SCC1-PK9::KAN MX rad61Δ::HGH MX LEU2::GAL1P-SIC1(9M)/HIS3P-GAL1/HIS3P-GAL2/GAL1P-GAL4 (single copy)* | This study | **K20891** |
| *S. cerevisiae* Mat a MATa *SCC1-PK9::KanMX rad61Δ::HGH MX smc3(R1008I)::HIS3 scc2-45::NAT MX (L545P D575G) LEU2::GAL1P-SIC1(9M)/HIS3P-GAL1/HIS3P-GAL2/GAL1P-GAL4(single copy)* | This study | **K22294** |
| *S. cerevisiae* Mat a MATa *SCC1-PK9::KanMX rad61Δ::HGH MX smc3(R1008I)::HIS3 LEU2::GAL1P-SIC1(9M)/HIS3P-GAL1/HIS3P-GAL2/GAL1P-GAL4(single copy)* | This Study | **K22296** |
| *S. cerevisiae* MATa *eco1Δ::NATMX4 rad61Δ::HGH MX scc2-45::NAT MX (L545P D575G) SMC3(E199A/R1008I)::HIS3 7.5 KB MINICHROMOSOME (TRP1 ARS1 CEN1-6KB)* | This study | **K22297** |
| *S. cerevisiae* MATa *eco1Δ::NATMX4 rad61Δ::HGH MX SMC3(E199A/R1008I)::HIS3 7.5 KB MINICHROMOSOME (TRP1 ARS1 CEN1-6KB)* | This study | **K22298** |
| *S. cerevisiae* MATa *SCC1(A547C)-PK6::KANMX SMC3(E570C,S1043C)::ADE2 SMC1(G22C,K639C)::NATMX 2.3 kb TRP1-ARS1-CEN4 plasmid* | Srinivasan et al. 2018 | **K23889** |
| *S. cerevisiae* MATa *SCC1(A547C)-PK6::KANMX SMC3(E570C,S1043C)::ADE2 SMC1(G22C,K639C)::NATMX LEU2::GAL1P-SIC1(9M)/HIS3P-GAL1/HIS3P-GAL2/GAL1P-GAL4 (single copy) 2.3 kb TRP1-ARS1-CEN4 plasmid* | Srinivasan et al. 2018 | **K23972** |
| *S. cerevisiae* MATa *SCC1(A547C)-PK6::KANMX SMC3(E570C,S1043C)::ADE2 SMC1(G22C,K639C)::NATMX scc2-45::NATMX (L545P D575G) 2.3 kb TRP1-ARS1-CEN4 plasmid* | Srinivasan et al. 2018 | **K24267** |
| *S. cerevisiae* MATa *SMC1(G22C,K639C)::NATMX4 SMC3(E570C,S1043C)::ADE2 LEU2::GAL-SCC1(R180E,R268D, A547C)-PK6::LEU2 SCC3 (K404E)-HA3::HIS 2.3 kb TRP1-ARS1-CEN4 plasmid* | This study | **K24697** |
| *S. cerevisiae* MATa *SCC1(A547C)-PK6::KANMX SMC3(E570C,S1043C)::ADE2 SMC1(G22C,K639C)::NATMX scc2-45::NATMX (L545P D575G) scc3 (K404E)-HA3::HIS 2.3 kb TRP1-ARS1-CEN4 plasmid* | This study | **K24709** |
| *S. cerevisiae* MATa *SMC1(G22C,K639C)::NATMX4 SMC3(E570C,S1043C)::ADE2 LEU2::GAL-SCC1(R180E,R268D, A547C)-PK6::LEU2 scc3 (K404E)-HA3::HIS scc2-45::natMX (L545P D575G) TRP1-ARS1-CEN4 2.3KB plasmid* | This study | **K24738** |
| *S. cerevisiae* MATa *trp1::smc3(E1155Q)-PK6::TRP1 smc1(D1164E)* | This study | **K25039** |
| *S. cerevisiae* MATa *SCC1(A547C)-PK6::KANMX SMC3(E570C,S1043C)::ADE2 SMC1(G22C,K639C)::NATMX scc2-45::NATMX (L545P D575G) 2.3kb Trp1-ARS1-Cen4 plasmid* | This study | **K25238** |
| *S. cerevisiae* MATa SCC1(A547C)-PK6::KANMX *pds5* (S81R)::HIS3 SMC3(E570C,S1043C)::ADE2 SMC1(G22C,K639C)::NATMX LEU2::GAL1P-SIC1(9M)/HIS3P-GAL1/HIS3P-GAL2/GAL1P-GAL4 (single copy) 2.3 kb TRP1-ARS1-CEN4 plasmid | This study | **K25311** |
| *S. cerevisiae* MATa SCC1(A547C)-PK6::KANMX *pds5* (S81R)::HIS3 SMC3(E570C,S1043C)::ADE2 SMC1(G22C,K639C)::NATMX *scc2-45::NATMX* (L545P D575G) LEU2::GAL1P-SIC1(9M)/HIS3P-GAL1/HIS3P-GAL2/GAL1P-GAL4 (single copy) 2.3 kb TRP1-ARS1-CEN4 plasmid | This study | **K25312** |
| *S. cerevisiae* MATa SCC1(A547C)-PK6::KANMX *scc3* (K404E)-HA3::HIS3 SMC3(E570C,S1043C)::ADE2 SMC1(G22C,K639C)::NATMX LEU2::GAL1P-SIC1(9M)/HIS3P-GAL1/HIS3P-GAL2/GAL1P-GAL4 (single copy) 2.3 kb TRP1-ARS1-CEN4 plasmid | This study | **K25313** |
| *S. cerevisiae* MATa SCC1(A547C)-PK6::KANMX *scc3* (K404E)-HA3::HIS3 SMC3(E570C,S1043C)::ADE2 SMC1(G22C,K639C)::NATMX *scc2-45::NATMX* (L545P D575G) LEU2::GAL1P-SIC1(9M)/HIS3P-GAL1/HIS3P-GAL2/GAL1P-GAL4 (single copy) 2.3 kb TRP1-ARS1-CEN4 plasmid | This study | **K25316** |
| *S. cerevisiae* MATa SCC1(A547C)-PK6::KANMX SMC3(E570C,S1043C)::ADE2 SMC1(G22C,K639C)::NATMX *scc2-45::NATMX* (L545P D575G) *cdc28-as1* | This study | **K25423** |
| *S. cerevisiae* MATa SMC3(E570C,S1043C)::ADE2 SMC1(G22C,K639C)::NATMX *leu2::Gal-Scc1*(R180E,R268D, A547C)-PK6::LEU2 *scc3* (K404E)-HA3::HIS3 *cdc28-as1* 2.3 kb TRP1-ARS1-CEN4 plasmid | This study | **K25437** |
| *S. cerevisiae* MATa SMC3(E570C,S1043C)::ADE2 SMC1(G22C,K639C)::NATMX *leu2::Gal-Scc1*(R180E,R268D, A547C)-PK6::LEU2 *scc2-45::NATMX* (L545P D575G) *scc3* (K404E)-HA3::HIS3 *cdc28-as1* 2.3 kb TRP1-ARS1-CEN4 plasmid | This study | **K25440** |
| *S. cerevisiae* MATa SMC3(E570C,S1043C)::ADE2 SMC1(G22C,K639C)::NATMX SCC1(A547C)-PK6::KANMX *scc2-45::natMX* (L545P D575G) *eco1Δ::HGH MX scc3* (K404E)-HA3::HIS TRP1-ARS1-CEN4 2.3KB plasmid | This study | **K25947** |
| *S. cerevisiae* MATa *ura::ADH1 promoter-OsTIR1-9myc::URA3 PDS5-Pk3-aid::KanMX4 scc2-45::NatMX scc3* (K404E)::HA3::HIS *leu2::Gal1p-Sic1*(9m)/His3p-Gal1/His3p-Gal2/Gal1p-Gal4::Leu2 (single copy) | This study | **K26414** |
| *S. cerevisiae* MATa *ura::ADH1 promoter-OsTIR1-9myc::URA3 PDS5-Pk3-aid::KanMX4 scc3* (K404E)::HA3::HIS *leu2::Gal1p-Sic1*(9m)/His3p-Gal1/His3p-Gal2/Gal1p-Gal4::Leu2 (single copy) | This study | **K26415** |
| *S. cerevisiae* MATa *smc1*(D1164E)::HIS3MX SCC1-PK9::KanMX *leu2::Gal1p-Sic1*(9m)/His3p-Gal1/His3p-Gal2/Gal1p-Gal4::Leu2 (single copy) | This study | **K26765** |
| *S. cerevisiae* MATa *smc1*(D1164E)::HIS3MX SCC1-PK9::KanMX *scc2-45::natMX* (L545P D575G) *leu2::Gal1p-Sic1*(9m)/His3p-Gal1/His3p-Gal2/Gal1p-Gal4::Leu2 (single copy) | This study | **K26766** |
| *S. cerevisiae* MATa *ura::ADH1 promoter-OsTIR1-9myc::URA3 scc2-3XmAID::KANMX6 Scc1-PK9::KanMX rad61Δ::HGH MX leu2::Gal1p-Sic1*(9m)/His3p-Gal1/His3p-Gal2/Gal1p-Gal4::Leu2 (single copy) | This study | **K26831** |
| *S. cerevisiae* MATa *scc2-45::natMX (L545P D575G)*<br>*leu2::Gal1p-Sic1(9m)/His3p-Gal1/His3p-Gal2/Gal1p-Gal4::Leu2 (single copy)* *ura3::ScclP-Smc3(E570C)-TEV3-Sccl(A547C)-PK9::KanMX::URA3 (single integrant,fusion linker: (GGGGS)x8+TEV3)* | This study | **K26993** |
| *S. cerevisiae* MATa *scc2-45::natMX (L545P D575G)*<br>*leu2::Gal1p-Sic1(9m)/His3p-Gal1/His3p-Gal2/Gal1p-Gal4::Leu2 (single copy)* *Smc1(G22C, K639C)::NAT MX*<br>*ura3::ScclP-Smc3(E570C)-TEV3-Sccl(A547C)-PK9::KanMX::URA3 (single integrant,fusion linker: (GGGGS)x8+TEV3)* | This study | **K26994** |
| *S. cerevisiae* MATa *bar1::hisG ura3-1::GAL-OsTIR1-9Myc::URA3 cdc45-AID (CDC45-5xGA-IAA17)::kanMX*<br>*rad61-3slAA::natNT2 SCC1-PK6::TRP1 scc2-45::natMX (L545P D575G)* | This study | **K27168** |
| *S. cerevisiae* MATa *bar1::hisG ura3-1::GAL-OsTIR1-9Myc::URA3 cdc45-AID (CDC45-5xGA-IAA17)::kanMX*<br>*rad61-3slAA::natNT2 SCC1-PK6::TRP1* | This study | **K27169** |
| *S. cerevisiae* MATa *TET R-GFP::LEU2*<br>*TETOs::URA3 pds5 S81R-HA6::HIS (NO HA tag)*<br>*trp1::MET3p-CDC20::TRP1* | This study | **K27443** |
| *S. cerevisiae* MATa *SCC1-PK9::KanMX smc3(T1185M)-HIS3MX6 scc2-45::natMX (L545P D575G)*<br>*leu2::Gal1p-Sic1(9m)/His3p-Gal1/His3p-Gal2/Gal1p-Gal4::Leu2 (single copy)* | This study | **K27536** |
| *S. cerevisiae* MATa *SCC1-PK9::KanMX smc3(T1185M)-HIS3MX6 leu2::Gal1p-Sic1(9m)/His3p-Gal1/His3p-Gal2/Gal1p-Gal4::Leu2 (single copy)* | This study | **K27537** |
| *S. cerevisiae* MATa *SCC1-PK9::KanMX*<br>*rad61Δ::hghMX leu2::Gal1p-Sic1(9m)/His3p-Gal1/His3p-Gal2/Gal1p-Gal4::Leu2 (single copy)* | This study | **K27569** |
| *S. cerevisiae* MATa *SCC1-PK9::KanMX*<br>*rad61Δ::hghMX scc4-2::NAT leu2::Gal1p-Sic1(9m)/His3p-Gal1/His3p-Gal2/Gal1p-Gal4::Leu2 (single copy)* | This study | **K27570** |
| *S. cerevisiae* MATa *scc2-4 TET R-GFP::LEU2*<br>*TETOs::URA3 pds5 S81R-HA6::HIS (NO HA tag)*<br>*trp1::MET3p-CDC20::TRP1* | This study | **K27575** |

